# A new double reporter strategy reveals a subset of non-migratory hematopoietic stem cells localized in a dynamic bone marrow niche

**DOI:** 10.1101/2025.07.20.665772

**Authors:** Marine A. Secchi, Qi Liu, Sara González Antón, Cera Mai, Aisha Jibril, Jiarui Xu, Dónal O’Carroll, Nicola K. Wilson, Kamil R. Kranc, Berthold Göttgens, Cristina Lo Celso, Tiago C. Luis

## Abstract

Interactions of hematopoietic stem cells (HSCs) with the bone marrow microenvironment are critical to regulate stem cell function in homeostasis, emergency hematopoiesis and ageing. The dynamic behavior of endogenous HSCs in their niches has been challenging to study due to the complexity of markers required to define pure HSCs. Multiple recently developed reporter strategies advanced our capacity to identify HSCs *in situ*. Yet, they provide different levels of HSC enrichment and are frequently not stable in situations of perturbed hematopoiesis, leading to contradictory observations of HSC migratory behavior. Here we employed a new double reporter strategy by combining the *Hoxb5*-mKO2 and *Vwf*-GFP reporters, that are stably expressed in homeostasis, stress and ageing. Hoxb5^+^Vwf^+^ cells represent a subset of highly pure and potent long-term HSCs with a platelet-biased pattern of differentiation, that are included in the populations identified by other reporter strategies. Intravital microscopy revealed Hoxb5^+^Vwf^+^ cells to be non-migratory in homeostasis, following platelet depletion when these cells are actively proliferating, and during ageing. This non-migratory behavior of HSCs in homeostasis and stress indicates that their activation does not inherently depend on relocation to alternative niches eliciting proliferation, as previously proposed. We found Hoxb5^+^Vwf^+^ HSCs in direct contact with vasculature and LepR^+^ perivascular cells but not preferentially closer to megakaryocytes. Nevertheless, increased megakaryopoiesis following platelet depletion brings megakaryocytes into close proximity of Hoxb5^+^Vwf^+^ HSCs, revealing previously unrecognized niche dynamics in HSC regulation during regeneration.

**DATA SHARING STATEMENT:** Original data will be made available upon reasonable request. The RNA sequencing data used in this study has been published before^35^ and is available from GEO (GSE81682). The code used for analysis is available from https://github.com/loversaber/Secchi_Paper_RNAseq_Analysis.git.

**KEY POINTS:**

1. Combined expression of *Hoxb5*-mKO2 and *Vwf*-GFP reporters defines a population of highly enriched long-term HSCs *in situ* and *in vivo*.
2. Hoxb5^+^Vwf^+^ HSCs are non-migratory in homeostasis, stress hematopoiesis and ageing, while their niche is highly dynamic.

## INTRODUCTION

Hematopoietic stem cells (HSCs) sustain homeostatic levels of blood cells throughout life thanks to their self-renewal and multilineage differentiation capacity. Additionally, they drive the regeneration of the hematopoietic system following acute stress conditions such as infection and bleeding, or clinical interventions like chemotherapy and bone marrow (BM) transplantation^1^. The BM microenvironments, or niches, occupied by HSCs play a critical role in regulating their key fate choices, including the decisions between remaining quiescent or initiating proliferation, and whether to self-renew or differentiate. Within the BM HSCs have been found in distinct anatomic regions and in close proximity to different stromal, endothelial and mature hematopoietic cell types, leading to the hypothesis that distinct HSC niches may exist^2,3^. Yet, the precise cellular composition of these niches and how HSCs interact with them are still highly controversial^4^, likely owning to the different levels of HSC enrichment achieved with the multiple phenotypic definitions employed to investigate HSCs *in situ*, and an overall low resolution in the identification of highly heterogeneous niche cell compartments^5-7^.

This led to a large effort in generating new reporter strategies to better identify HSCs *in situ* and *in vivo*^8-16^. Ideally, such reporters will be specifically and uniformly expressed in HSCs, with specificity to only an HSC subset being preferable to broader expression that includes other cell types. They should be stable in homeostasis, ageing and in different scenarios of perturbed hematopoiesis, whilst also sufficient for HSC identification, circumventing the need for additional cell surface markers. This allows bypassing the requirement for transplantation and associated BM conditioning, thereby enabling longitudinal studies of the HSCs’ dynamic interactions with the native microenvironment in homeostasis and stress, which is made possible by intravital microscopy (IVM)^17-20^. Among these strategies, *Hoxb5* reporters^9,12^ confer a high level of HSC enrichment, even though some very low long-term (LT) engraftment has been detected within the Hoxb5^-^ HSC compartment, and Hoxb5^+^ cells were shown to have a near homogenous perivascular localization in BM^9^. More recently, 3 new *in vivo* reporter strategies leading to high levels of HSC enrichment within the population of reporter-positive cells revealed contrasting behaviors of endogenous HSCs in homeostasis and following mobilization treatments. Using a *Mds1*-GFP reporter transgene that becomes inactivated following *Flt3*-Cre mediated recombination (MFG reporter), Christodoulou *et al*. reported low HSC motility in homeostasis, and a heterogenous, dynamic behavior following treatment with cyclophosphamide and G-CSF that leads to expansion and mobilization of HSCs^10^. This was in line with the stationary behavior of *Hlf-*TdTom^hi^ cells in adult mice reported by Takihara *et al.*^15^. Conversely, using a fate-mapping reporter strategy where HSCs are labelled following tamoxifen-induced *Pdzk1ip1*-CreER mediated recombination, Upadhaya *et al*. reported significant HSC motility, which is rapidly abrogated in response to mobilization induced through inhibition of Cxcr4 and integrin signalling^16^. While these differences might be explained by the different transgenic strategies employed labelling distinct HSC subsets, they highlight the need for new models to further investigate the behavior of endogenous HSCs *in vivo*.

The HSC compartment is highly heterogeneous, comprising different subsets of lineage-biased HSCs^21-23^, and including a platelet-biased subset identified by single cell transplantations and *in steady state by in vivo* genetic barcoding assays^14,24-27^. Platelet-biased HSCs express the von Willebrand factor (*Vwf*) gene allowing their prospective identification *in vivo*, using *Vwf* reporter transgenes^26,28,29^. Importantly, we recently showed that Vwf^+^ HSCs are preferentially recruited into proliferation following acute platelet depletion, with the majority of Vwf^+^ HSCs transitioning into S-G2-M phase of cell cycle within 24 hrs^26,29^. This is part of a feedback mechanism that contributes to reestablishing platelet homeostasis and that depends on IL-1 signaling in LepR^+^ perivascular niche cells^29^. Megakaryocytes (MKs), have been implicated as critical regulators of HSC quiescence^30,31^ and, importantly, Vwf^+^ HSCs have been suggested to occupy a distinct niche, closer to MKs and farther from arterioles in BM^32^. Distinct niche occupancy for quiescent and activated HSCs has been proposed, implying a dynamic behavior where HSCs may migrate from quiescence reinforcing niches to alternative environments eliciting proliferation^4,8,10,33,34^. The specific response of Vwf^+^ HSCs to platelet depletion offers the unique opportunity to test if their preferential activation results from changes in HSC-niche interactions, due to HSC relocation or changes in niche composition.

Here we employed a new double reporter strategy by combining the *Hoxb5*-mKO2^12^ and the *Vwf*-GFP^26^ reporters to identify highly enriched HSCs *in situ* and *in vivo,* and longitudinally investigate their behavior in homeostasis, stress and ageing. IVM revealed Vwf^+^ HSCs to be highly confined within the BM niche in homeostasis, following platelet depletion and during ageing, demonstrating that relocation to a separate niche is not essential for activation of HSC proliferation. Yet, highly dynamic megakaryopoiesis leads to increased proximity between MKs and Vwf^+^ HSCs following platelet depletion, unravelling a novel aspect of BM niche dynamics in the regulation of HSCs under stress conditions.

## RESULTS

### Combined expression of *Hoxb*5-mKO2 and *Vwf*-GFP defines a highly enriched LT-HSC population

To increase the purity of reporter-labeled HSCs compared to existing strategies and achieve greater precision in tracking HSCs by IVM, we crossed *Hoxb5*-mKO2^12^ with *Vwf*-GFP^26^ transgenic reporter mice and analyzed the expression of Kusabira Orange (mKO2) and GFP in HSCs. The expression of *Hoxb*5-mKO2 and *Vwf*-GFP reporter transgenes within Lin^-^Sca1^+^cKit^+^CD48^-^ CD150^+^ (LSK-SLAM) HSCs of young adult mice in homeostasis defines 3 main subsets of cells, with 17.5±5.2% of cells co-expressing both reporters (LSK-SLAM Hoxb5^+^Vwf^+^), 18.3±2.1% of cells expressing only *Hoxb5*-mKO2 (LSK-SLAM Hoxb5^+^Vwf^-^), and 60.2±6.5% of cells expressing neither (LSK-SLAM Hoxb5^-^Vwf^-^) (**Fig.1A-B**). In line with previous studies demonstrating that the majority of HSC activity is present within the Hoxb5^+^ population^9,12^, cells in the Hoxb5^+^Vwf^+^ and Hoxb5^+^Vwf^-^ LSK-SLAM subsets express high levels of EPCR and are mostly negative for CD34 expression, while the Hoxb5^-^Vwf^-^ population exhibits significantly lower levels of EPCR expression and frequency of CD34^-^ cells (**Fig.1C-D**).

**Figure 1.**
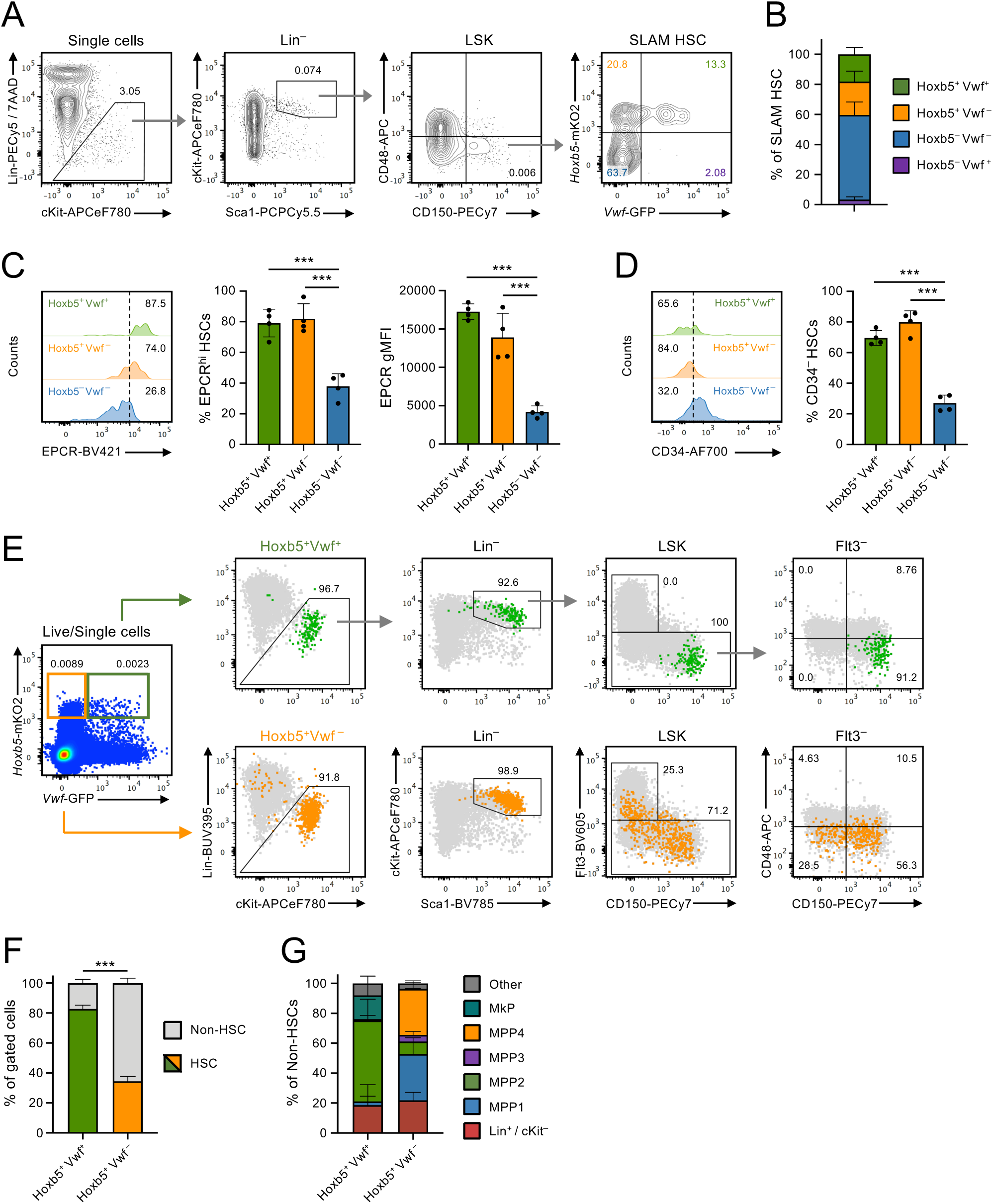
Combined expression of *Hoxb*5-mKO2 and *Vwf*-GFP defines highly enriched phenotypic HSC populations. **A-B)** Flow cytometric analysis of *Hoxb5*-mKO2 and *Vwf*-GFP reporters’ expression in phenotypically defined SLAM-HSCs. **A)** Representative gating strategy of SLAM-HSCs with frequency of the subsets defined by *Hoxb5*-mKO2 and *Vwf*-GFP expression. Numbers in gates/quadrants indicate the frequency of the gated cell populations among single cells, except for last plot (right) where they represent frequencies of parent SLAM-HSC population, for the representative mouse shown. Data in (**B**) are mean±SD frequencies from 4 mice in 3 independent experiments. **C-D)** Flow cytometric analysis of EPCR **(C)** and CD34 **(D)** expression within the SLAM-HSC subsets defined by *Hoxb5*-mKO2 and *Vwf*-GFP expression. Histograms show 1 representative mouse of a total of 4 presented in graphs. Numbers in histograms indicate the frequency of EPCR^hi^ **(C)** and CD34^-^ cells **(D)** within the indicated subsets. Data from 3 independent experiments. gMFI, geometric mean fluorescence intensity. ***, p<0.001 using 1-way ANOVA with Tukey’s multiple comparisons test. **E-G)** Phenotypic characterization of total Hoxb5^+^Vwf^+^ and Hoxb5^+^Vwf^-^ cells in BM, defined exclusively based on the expression of the reporter transgenes. **E)** Hoxb5^+^Vwf^+^ (green) and Hoxb5^+^Vwf^-^ (orange) cells were gated from total BM live single cells and analysed for the frequency of SLAM-HSCs. Numbers next to gates/quadrants indicate frequency of the gated cells among the parent population. Grey cells in the background represent total BM cells undergoing same gating strategy. **F)** Mean±SD frequency of SLAM-HSCs within the Hoxb5^+^Vwf^+^ and Hoxb5^+^Vwf^-^ populations. ***, p<0.001 (t-test, for the frequency of HSCs). **G)** Phenotypic analysis of the populations represented within the non-HSC fractions of the total Hoxb5^+^Vwf^+^ and Hoxb5^+^Vwf^-^ cells in BM. “Other” include cells outside defined gates and populations with <0.5% representation (including pre-MegE, pre-CFU-E, pre-GM and GMP). Data are mean±SD frequencies from 8 mice in 4 independent experiments. See also Supplementary Figure 1.

We subsequently analyzed the frequency of phenotypic HSCs (LSK-SLAM) within the populations solely defined by the expression of *Hoxb5* and *Vwf* reporters (**Fig.1E**). Remarkably, combined expression of *Hoxb*5-mKO2 and *Vwf*-GFP defines an extremely rare population representing 0.0017±0.0006% of total live cells in BM that is highly enriched for phenotypically defined HSCs (83±2.5%; **Fig.1F**) and homogeneously express high levels of the SLAM HSC marker CD150 (**Fig.1E**). The minor non-HSC fraction of the total Hoxb5^+^Vwf^+^ population (17±2.5%) is mostly constituted by LSK Flt3^-^CD48^+^CD150^+^ megakaryocyte-primed multipotent progenitors 2 (MPP2), Lin^-^Kit^+^Sca1^-^CD41^+^CD150^+^ megakaryocyte progenitors (MkP) and Lin^+^/cKit^-^ cells (**Fig.1E,G**; **Supplementary** Fig.1A). Conversely, the Hoxb5^+^Vwf^-^ cells constitute a more abundant (0.0055±0.0017%) and more heterogeneous population constituted of 34±3.3% LSK-SLAM cells (**Fig.1E-F**) and, within the non-HSC fraction, of LSK Flt3^-^CD48^-^CD150^-^ MPP1 (or short-term HSCs; ST-HSCs), lymphoid-primed LSK Flt3^+^ MPP4 and Lin^+^/cKit^-^ cells in approximately similar proportions. (**Fig.1G**).

To obtain further evidence at transcriptional level for differences in stemness between Hoxb5^+^Vwf^+^ and Hoxb5^+^Vwf^-^ cell populations identified by the expression of the 2 reporters alone we next analyzed a previous single-cell RNA-sequencing (scRNA-Seq) data set performed on total hematopoietic stem and progenitor cells (HSPC; LSK cells)^35^ Cells expressing *Hoxb5* or *Vwf* were identified based on transcript levels across the entire data set (**Supplementary** Fig.1B). Co-expression of both genes identified a small population of cells mostly confined within the LT-HSC population, while cells expressing only *Hoxb5* were more dispersed throughout the HSPC landscape (**Fig.2A**). Of note, using the previously developed HSC score^36,37^ and Repopulation score (RepopSig)^38^, we observed increased expression of these transcriptional signatures in *Hoxb5*^+^*Vwf*^+^ cells, when compared with *Hoxb5*^+^*Vwf*^-^ cells or even to the total LT-HSC population, suggesting higher HSC function in the *Hoxb5*^+^*Vwf*^+^ population (**Fig.2B-C**). Since by flow cytometry the Hoxb5^+^Vwf^+^ cells only represented 17.5% of the total LSK-SLAM population (**Fig.1B**) we further interrogated the Nestorowa *et al.* dataset to compare the overlap between *Hoxb5*^+^*Vwf*^+^ cells and other reporter-based definitions used to identify HSCs *in situ* and *in vivo.* We identified *Mds1^+^Flt3^-^*, *Tcf15^+^*, *Pdzk1p1^+^* and *Ctnnal1^+^Hlf^+^* cells^8,10,14-16^ (**Fig.2D**) based on transcript levels of these genes across the data set (**Supplementary** Fig.1C). To better reflect the expression of the MFG (*Mds1^+^Flt3^-^*) reporter^10^, expression of *Mds1* was analyzed by quantifying reads specifically within the first 2 *Mds1* exons, rather than the whole *Mecom* complex locus, that also includes *Evi1* and is more broadly expressed throughout the HSPC landscape (**Supplementary** Fig.1D**-E**). *Hlf* was used in combination with *Ctnnal1* as a proxy for *Kit* expression. Of note, 96% of *Hoxb5^+^Vwf^+^* cells overlapped with cells identified by the other HSC definitions, although they only constitute a minor fraction of the populations identified by these other strategies (10.6% of *Mds1^+^Flt3^-^* cells, 9.5% of *Pdzk1p1^+^*cells, 7.4% of *Tcf15^+^* cells and 14.3% of *Ctnnal1^+^Hlf^+^*cells) (**Fig.2E-H**). Furthermore, approximately 50% of *Mds1^+^Flt3^-^* and *Pdzk1p1^+^*cells, and 40% of *Tcf15^+^* and *Ctnnal1^+^Hlf^+^*cells were not overlapping with any other identification strategy, and importantly, we did not find any cell in common between all 5 HSC definitions analyzed (**Fig.2E-H**). In line with the lower enrichment of LSK-SLAM cells within the Hoxb5*^+^*Vwf^-^ population defined by reporters’ expression (**Fig.1F**), 40% of *Hoxb5^+^Vwf^-^* cells did not overlap with any other HSC definition (**Fig.2F, H**). Together, this analysis demonstrates a high level of heterogeneity in the cells identified by different reporter strategies. Common to all strategies is the fact that each identifies only a subset of total HSCs. The new Hoxb5^+^Vwf^+^ definition identifies the totality of the Vwf^+^ HSC subset (**Fig.1B**) that has been functionally linked with platelet-MK differentiation bias^24,26^, and these cells express combinations of all other reporters used, rather than constituting their own HSC subset. Thus, this dual reporter strategy allows easy identification and purification of HSCs expressing a core HSC signature, akin to the MolO HSCs previously identifiable solely by transcriptomic analyses^36,37^.

**Figure 2.**
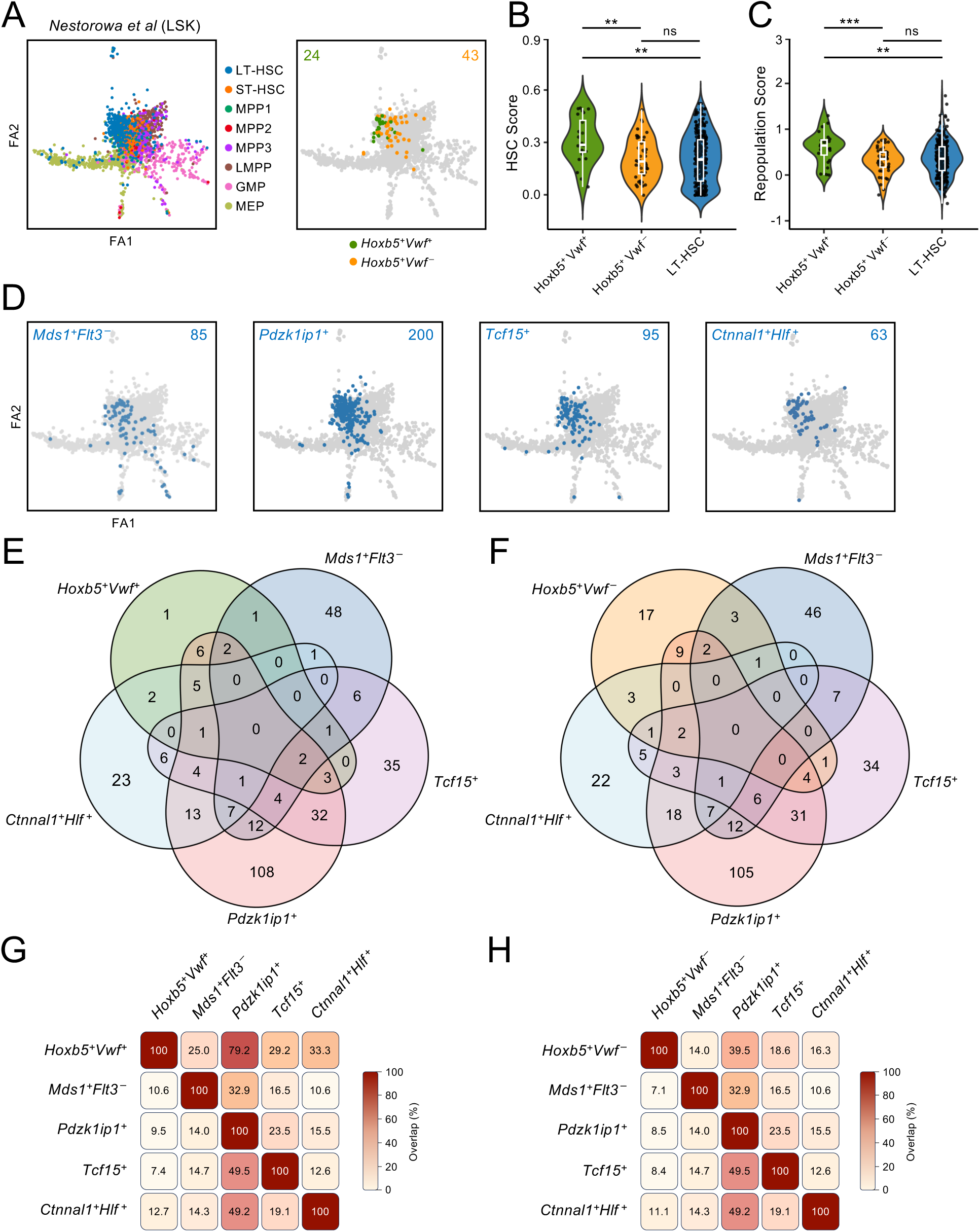
Hoxb5^+^Vwf^+^ HSCs are a subset of partially overlapping alternative HSC reporter strategies. **A)** Single cell RNA sequencing analysis of total LSK cells from the Nestorowa *et al.*^35^ data set showing a force directed graph of the landscape with highlighted immunophenotypic populations (left), and Hoxb5^+^Vwf^+^ and Hoxb5^+^Vwf^-^ cells identified based on *Hoxb5* and *Vwf* transcript levels (right). **B-C)** HSC score **(B)** and Repopulation score **(C)** of Hoxb5^+^Vwf^+^ cells, Hoxb5^+^Vwf^-^ cells, and total LT-HSCs identified based on immunophenotypic definition from index sorting. *Vwf* transcript levels were excluded from score calculations as this gene is part of the definition used here to identify the cells. **D)** Projections of *Mds*1^+^*Flt3*^-^, *Pdzk1ip1*^+^, *Tcf15*^+^ and *Ctnnal1*^+^*Hlf*^+^ cells, identified based on transcript levels, on the Nestorowa LSK landscape. Numbers in graph indicate the number of cells identified using each HSC definition presented. **E-F)** Venn diagrams depicting the number of Hoxb5^+^Vwf^+^ **(E)** and Hoxb5^+^Vwf^-^ cells **(F)** cells overlapping with each HSC definition used for comparison. **G-H)** Frequency of overlapping cells between all HSC definitions analysed, showing comparisons with Hoxb5^+^Vwf^+^ cells **(G)** and Hoxb5^+^Vwf^-^ cells **(H)**. See also Supplementary Figure 1.

To confirm the high LT-HSC activity of Hoxb5^+^Vwf^+^ cells, we competitively transplanted limiting numbers of double and single positive cells (CD45.2) isolated based on reporters’ expression alone (**Supplementary** Fig.2A), into lethally irradiated mice (CD45.1). Hoxb5^+^Vwf^+^ cells revealed a 1/5.8 frequency of reconstitution units (**Fig.3A**). In line with *Vwf*-GFP expression defining a subset of platelet-biased HSCs^24,26^, Hoxb5^+^Vwf^+^ cells conferred robust LT multilineage reconstitution (**Fig.3B**, **Supplementary** Fig.2B**-C**) with a platelet-biased pattern, as evidenced by a lineage reconstitution Platelet/Myeloid ratio >1 and in comparison with the Hoxb5^+^Vwf^-^ counterparts (**Fig.3C**). While *in vivo* limiting dilution analysis of Hoxb5^+^Vwf^-^ cells revealed a non-statistically significant difference in the frequency of reconstitution units when compared to Hoxb5^+^Vwf^+^ cells (**Fig.3A**), the level of lineage reconstitution was significantly lower, particularly in the platelets and myeloid lineages (**Fig.3B**). Furthermore, Hoxb5^+^Vwf^+^ cells conferred a more robust and significantly higher reconstitution of the LSK-SLAM HSC compartment of primary recipient mice (**Fig.3D**) and were able to reconstitute all 3 subsets defined by *Hoxb*5-mKO2 and *Vwf*-GFP expression within the LSK-SLAM HSCs (**Fig.3E** and **Fig,1B**). By contrast, the Hoxb5^+^Vwf^-^ cells failed to reconstitute the Vwf^+^ HSC compartment, as previously reported^26^. Lastly, to confirm the high frequency of functional LT-HSCs within the Hoxb5^+^Vwf^+^ compartment, we performed secondary transplants. Whole BM from primary recipients transplanted with 10 Hoxb5^+^Vwf^+^ cells conferred high LT multilineage reconstitution of both peripheral blood and the HSC compartment in secondary recipient mice (**Fig.3F-G**). Contrarily, whole BM from primary recipients reconstituted with 10 Hoxb5^+^Vwf^-^ cells mostly failed to reconstitute secondary recipients. This is in agreement with the higher heterogeneity and higher frequency of MPP1/ST-HSCs and MMP4 cells observed in the Hoxb5^+^Vwf^-^ population (**Fig.1F-G**) and lower HSC and repopulation scores in these cells (**Fig.2B-C**). Together, these data demonstrated the suitability of the *Hoxb*5-mKO2 and *Vwf*-GFP dual reporter strategy to identify phenotypic and functionally defined Vwf^+^ HSCs endogenously and without the need of other markers.

**Figure 3.**
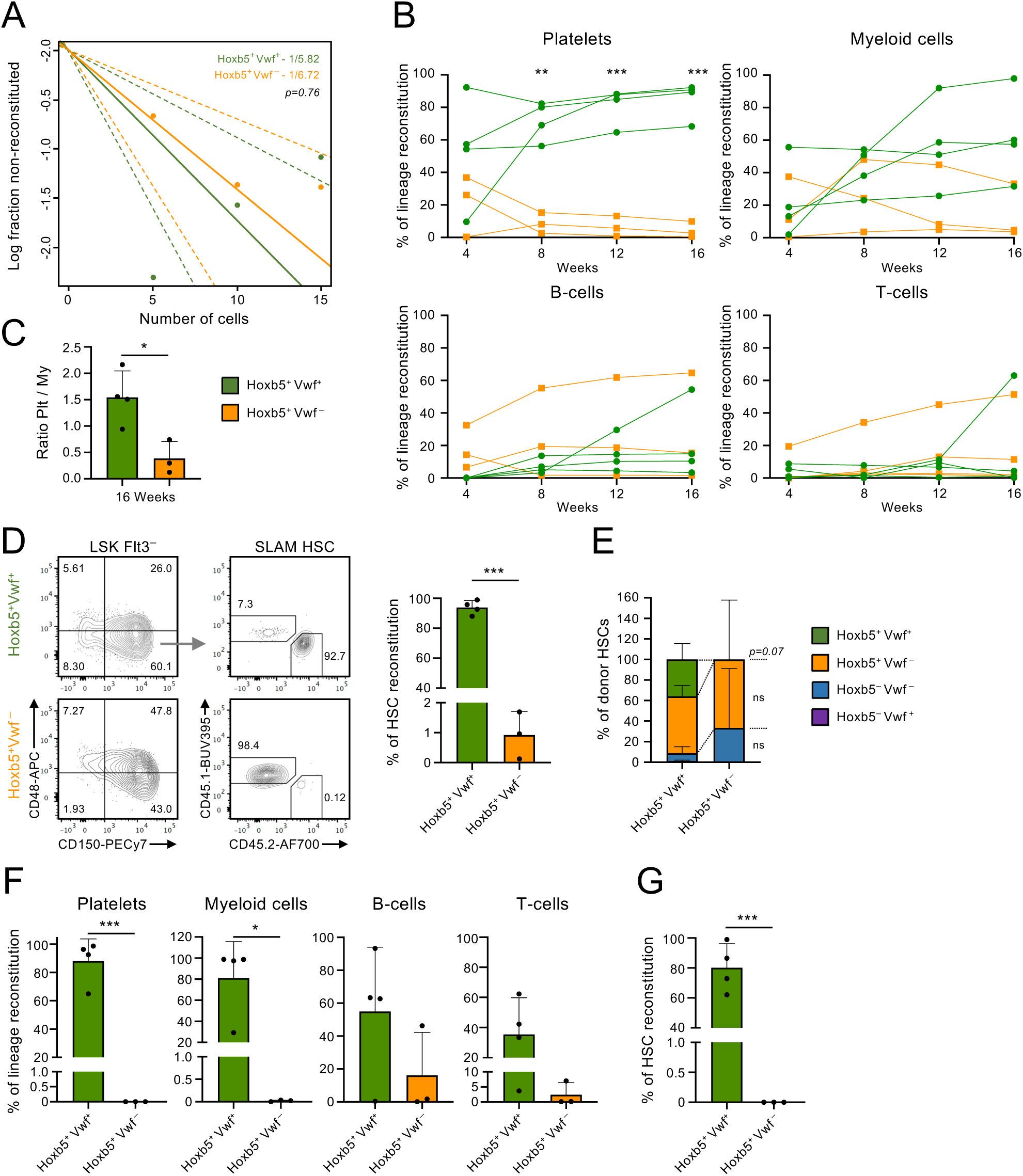
Combined expression of *Hoxb5*-mKO2 and *Vwf-*GFP in bone marrow defines highly functional long-term HSCs. **A)** Limiting dilution analysis of *in vivo* reconstitution capacity of Hoxb5^+^Vwf^+^ and Hoxb5^+^Vwf^-^ FACS sorted cells from total live BM cells, based only on the expression of the *Hoxb5*-mKO2 and *Vwf*-GFP reporter transgenes. Recipient mice were transplanted with 5, 10 or 15 Hoxb5^+^Vwf^+^ or Hoxb5^+^Vwf^-^ cells and analysed for multilineage long-term (16 weeks) reconstitution in peripheral blood. Data from a total of 13 and 12 mice transplanted with Hoxb5^+^Vwf^+^ and Hoxb5^+^Vwf^-^ cells respectively. Numbers inside plot indicate the frequency of reconstitution units and *p* value calculated with ELDA software. **B-E)** Long-term reconstitution analysis of the platelet, myeloid and lymphoid cell lineages in peripheral blood at the indicated time points **(B-C)** and of the HSC compartment in BM 16 weeks post transplantation **(D-E)** of mice (CD45.1) reconstituted with 10 Hoxb5^+^Vwf^+^ or Hoxb5^+^Vwf^-^ cells (CD45.2). In **(B)** each line represents an individual mouse. **C)** Ratio of Platelet/Myeloid reconstitution. **D)** Representative FACS plots (left) and quantification (right) of CD45.1/CD45.2 reconstitution in LKS SLAM HSCs. **E)** *Hoxb5*-mKO2 and *Vwf*-GFP subsets composition of the CD45.2 SLAM-HSC compartment of the reconstituted mice. **D-E)** Data shows mean±SD of 4 and 3 mice transplanted with Hoxb5^+^Vwf^+^ or Hoxb5^+^Vwf^-^ cells, respectively. Only long-term reconstituted mice were included in this analysis. **F-G)** Total BM from primary reconstituted mice was transplanted into secondary recipients (CD45.1) and analysed for multilineage reconstitution in peripheral blood **(F)** and for HSC reconstitution in BM **(G)**, 16 weeks post transplantation. *, p<0.05; **, p<0.01; ***, p<0.001; ns, non-significant (p>0.05); using 2-way ANOVA with Sydak’s multiple comparisons **(A, E)** or t-test **(C-D, F-G)**. See also Supplementary Figure 2.

### Hoxb5^+^Vwf^+^ HSCs are sessile in homeostasis and after acute platelet depletion

To test whether HSC fate changes are linked to relocation to different niches, we monitored the behavior of Hoxb5^+^Vwf^+^ cells in the BM niche using calvarium IVM. Vwf^+^ HSCs have a platelet-biased pattern of lineage differentiation *in vivo* and are preferentially recruited into proliferation following acute platelet depletion^24-27,29^. Therefore, in addition to analyzing the dynamics of HSCs in homeostasis, we also investigated the behavior of these cells post platelet depletion (**Fig.4A**). We performed IVM immediately following anti-GPIbα antibody administration, when platelet activation and subsequent depletion are taking place (**Fig.4B**), and 24 hrs post platelet depletion, when the majority of Vwf^+^ HSCs have exited quiescence and are actively cycling^29^. Longitudinal IVM preformed prior to and immediately after anti-GPIbα antibody administration confirmed platelet depletion: in steady state platelets generated GFP^+^ signals within BM vessels, often distorted into lines because of the effect of blood flow and speed of confocal acquisition. However, this signal entirely disappeared within minutes post anti-GPIbα antibody administration (**Supplementary** Fig.3A). Flow cytometric characterization of the BM of double reporter mice 24 hrs post platelet depletion revealed no major changes in the SLAM-HSC compartment in terms of the composition of subsets defined by *Hoxb5*-mKO2 and *Vwf*-GFP expression. Apart from a reduction in the frequency of Hoxb5^+^Vwf^-^ cells and a corresponding increase in the frequency of Hoxb5^-^Vwf^-^ cells, there was no significant change in the frequency of Hoxb5^+^Vwf^+^ cells (**Supplementary** Fig.3B**-C**). The frequencies of EPCR^hi^ cells within LSK-SLAM Hoxb5^+^Vwf^+^ and Hoxb5^+^Vwf^-^ subsets also remained comparable (**Supplementary** Fig.3D**-E**). Finally, we confirmed similar enrichment of LSK-SLAM HSCs within the Hoxb5^+^Vwf^+^ and Hoxb5^+^Vwf^-^ cell populations defined exclusively based on the reporters’ expression in total live BM cells (**Supplementary** Fig.3F**-G**). The composition of the non-HSC fraction of the Hoxb5^+^Vwf^+^ population remained unchanged, while we observed a significant increase in MPP4 cells in the Hoxb5^+^Vwf^-^ non-HSC population (**Supplementary** Fig.3H).

**Figure 4.**
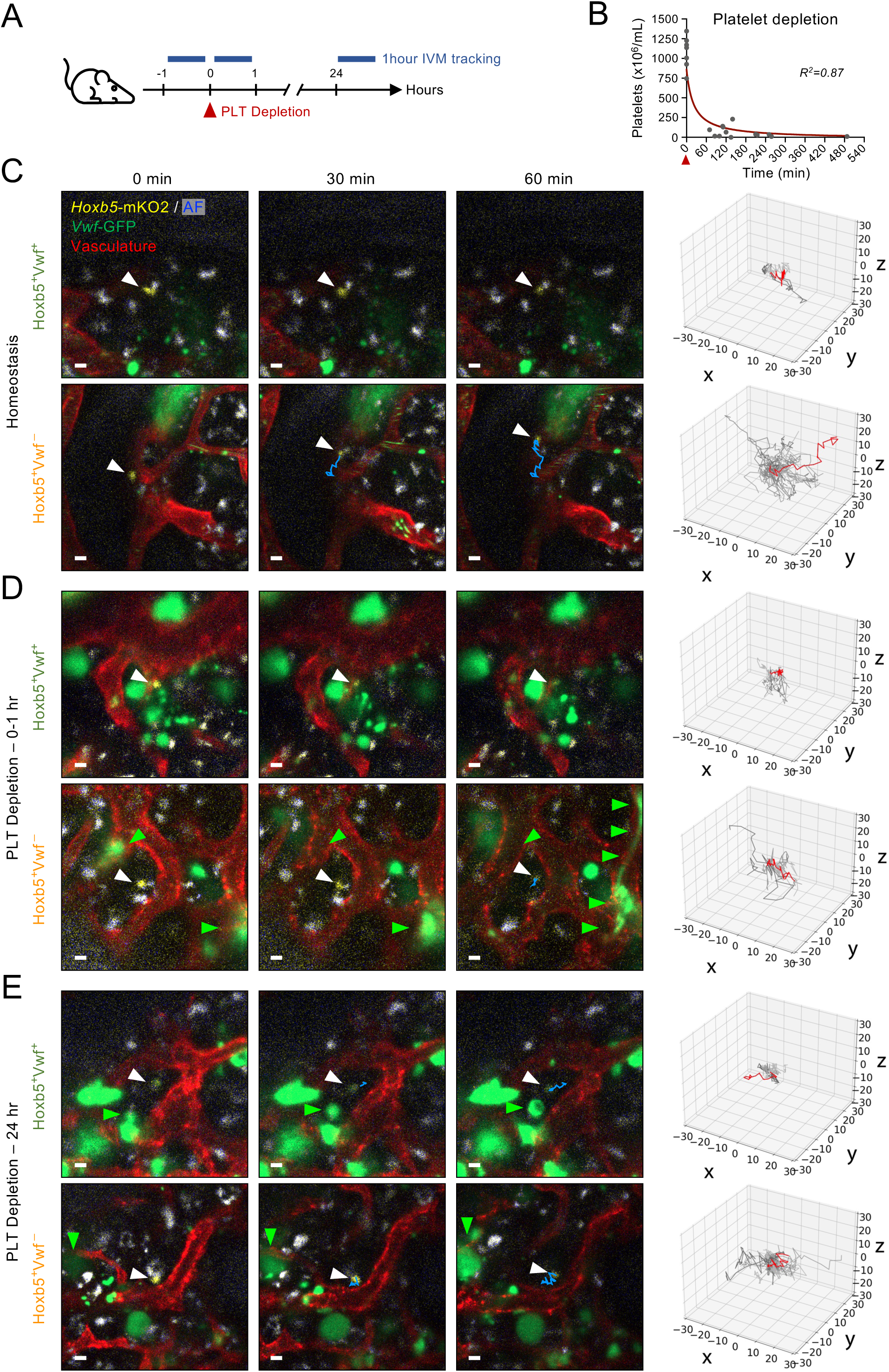
Intravital microscopy reveals very slow dynamics of Hoxb5^+^Vwf^+^ cells in homeostasis and post platelet depletion. **A)** The calvarium BM of double reporter mice was imaged by intravital microscopy (IVM) for 1 hr before platelet depletion, during the first hour and/or 24 hrs post administration of the anti-GPIbα antibody for platelet depletion. **B)** Non-linear fit curve of platelet numbers in peripheral blood before and during the first 8 hours post anti-GPIbα antibody administration. Data from 7 mice in homeostasis and 13 mice analysed at the indicated time-points. **C-E**) Selected timeframes at the indicated times, from time-lapse IVM, showing the migratory behaviours of representative Hoxb5^+^Vwf^+^ and Hoxb5^+^Vwf^-^ cells (arrowheads point at HSCs, blue lines represent the cells’ tracks) in the calvarium BM of mice in homeostasis **(C)**, and responding to platelet depletion (**D,** 0-1 hr; **E**, 24 hrs). Green arrowheads depict MKs growing, disappearing or forming pro-platelets during the 1 hr observation period. Scale bars represent 10 μm. Rose plots on the right show the 3-dimensional tracks (in μm) of all cells analysed (grey) in each category, with those from the cells depicted in the left images highlighted in red. Tracks for each cell were normalized for position of origin. N= 6, 8 and 9 Hoxb5^+^Vwf^+^ cells and 28, 10 and 28 Hoxb5^+^Vwf^-^ cells from mice in homeostasis, 0-1 hr, and 24 hrs post platelet depletion, respectively, from a total of 28 imaged mice. See also Supplementary Figures 3-4.

We then imaged Hoxb5^+^Vwf^+^ HSCs contained within frontal bone cavities in calvarium BM and compared them to Hoxb5^+^Vwf^-^ cells from the same mice (**Supplementary** Fig.4A). Hoxb5^+^ cells were identified by subtracting the autofluorescence from the mKO2 signal and subsequently assessed for GFP expression (**Supplementary** Fig.4B). In line with the low frequency of Hoxb5^+^Vwf^+^ cells in total BM and high enrichment for HSCs (**Fig.1E, Fig.2A**), these cells were extremely rare in the calvarium and we could typically not image more than 1 cell per mouse calvarium in homeostasis and immediately after depletion. Yet, in agreement with the strong cell cycle activation post platelet depletion, 2 and 3 Hoxb5^+^Vwf^+^ cells could be observed in the same field of view in 2 out of 6 mice analyzed 24 hrs post anti-GPIbα antibody administration (**Supplementary** Fig.4C). Remarkably, 3D time-lapse imaging of these cells during a 1 hr time window revealed that Hoxb5^+^Vwf^+^ HSCs homogenously exhibited a stationary behavior in homeostasis (**Fig.4C**) at both time points following platelet depletion (**Fig.4D-E; Supplementary Movies 1-6**), and only rarely moved somewhere else within the field of view (e.g. red track in **Fig.4E** top rose plot). Compatible with the higher cell heterogeneity found within the Hoxb5^+^Vwf^-^ population (**Fig.1E-G**), these cells revealed a more heterogeneous behavior, with the majority of the cells appearing migratory and moving within the field of view (**Fig.4C**). Supporting these observations, track analysis of individual Hoxb5^+^Vwf^+^ HSCs in homeostasis showed that these cells had no or very little maximum displacement, which measures the furthest distance from the starting point of each track (**Fig.5A**) and a very high arrest coefficient, measured as the percentage of time cells move less than 2 µm/min (**Fig.5B**). This confirmed that they remained within small, confined BM regions. Conversely, Hoxb5^+^Vwf^-^ cells covered significantly longer distances within the observation period and displayed a wider range of arrest coefficients, with a trend towards a higher proportion of cells displaying low arrest coefficient (**Fig.5A-B**). Interestingly, observation of timelapse data compiled into movie format clearly indicated that Hoxb5^+^Vwf^+^ cells were not entirely immobile and rather wiggled, engaging with the immediate microenvironment with dynamic and most often oscillatory interactions (**Supplementary Movies 1-6**). This type of oscillatory movement generates circulatory tracks without net migration. As a result, despite the significantly lower maximum displacement of the Hoxb5^+^Vwf^+^ cells, both Vwf^+^ and Vwf^-^ cell subsets displayed comparable total track lengths (**Fig.5C**) and mean speed (**Supplementary** Fig.5A). Consistent with this, Hoxb5^+^Vwf^+^ cells showed significantly reduced linearity coefficient (**Fig.5D**), which measures the directionality of the movement, and mean squared displacement (**Fig.5E**), reflecting a reduced diffusion, when compared to Hoxb5^+^Vwf^-^ cells. Lastly, both Hoxb5^+^Vwf^+^ and Hoxb5^+^Vwf^-^ cells showed a low and similar variance of speed, indicating a persistent behavior (**Fig.5F**). Remarkably, following platelet depletion the behavior of both Hoxb5^+^Vwf^+^ and Hoxb5^+^Vwf^-^ cells remained mostly unchanged, with neither population becoming more motile, which would be consistent with niche relocations. Instead, Hoxb5^+^Vwf^+^ cells exhibited reduced track length and mean speed immediately following depletion (**Fig.5C**, **Supplementary** Fig.5A), and Hoxb5^+^Vwf^-^ cells presented reduced maximum displacement, linearity coefficient and mean square displacement at 24 hrs post platelet depletion (**Fig.5A, D-E**). Altogether, this analysis revealed an unexpected non-migratory, yet oscillatory behavior of Hoxb5^+^Vwf^+^ HSCs and demonstrated that activation of Vwf^+^ HSCs following platelet depletion is not associated with their relocation to a different niche, unlinking changes in HSC cell cycle status with migration towards specialized niches that would support either HSC quiescence or proliferation.

**Figure 5.**
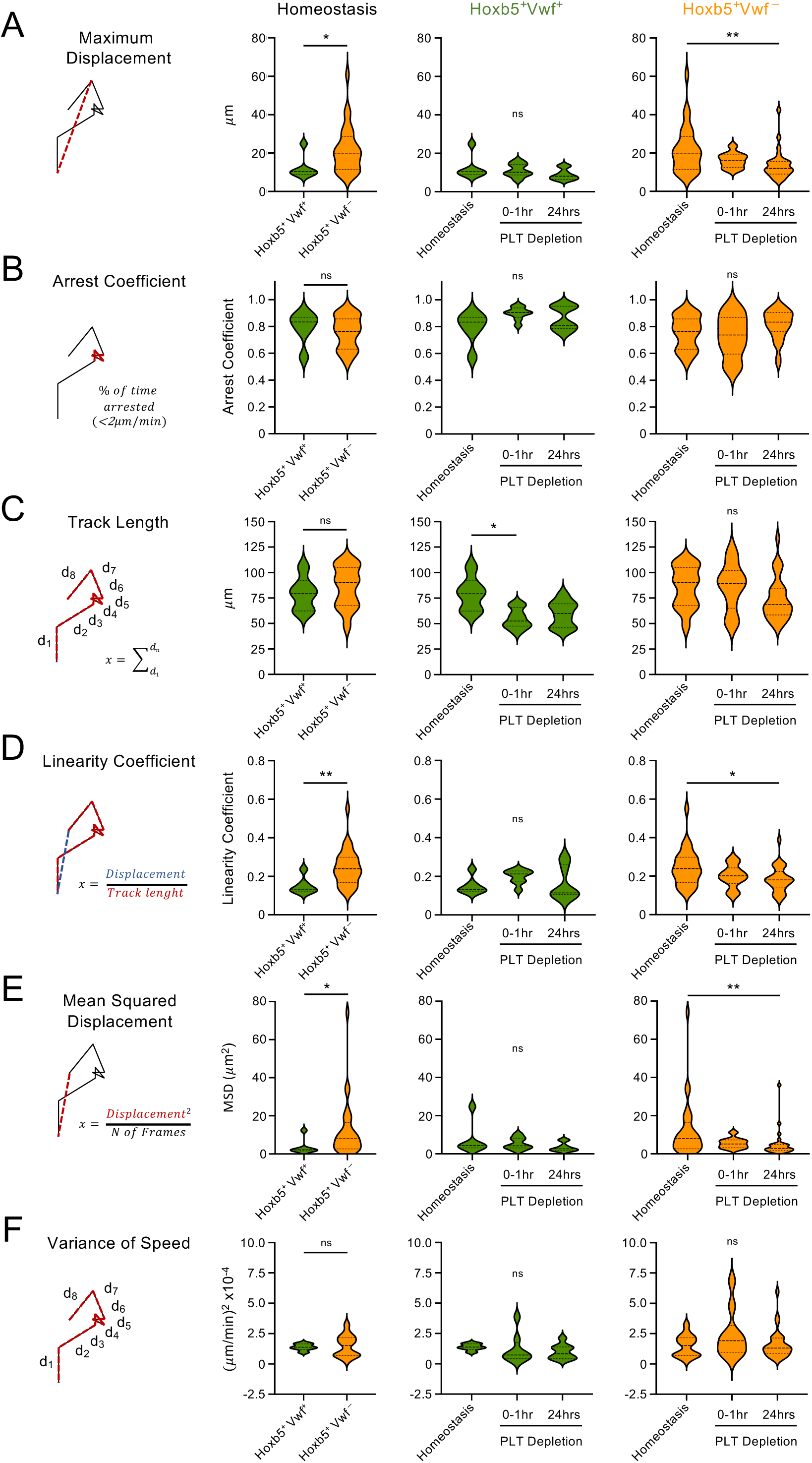
The Hoxb5^+^Vwf^+^ cells are not migratory in homeostasis and post platelet depletion. Track parameters extracted from the 3D time lapse IVM of single Hoxb5^+^Vwf^+^ and Hoxb5^+^Vwf^-^ cells (presented in Fig.4). Violin plots depict maximum displacement **(A)**, arrest coefficient **(B)**, track length **(C)**, linearity coefficient **(D)**, mean squared displacement **(E)** and variance of speed **(F)**. Data from 6, 8 and 9 Hoxb5^+^Vwf^+^ cells and 28, 10 and 28 Hoxb5^+^Vwf^-^ cells from mice in homeostasis, 0-1 hr, and 24 hrs post platelet depletion, respectively, from a total of 28 imaged mice. *, p<0.05; **, p<0.01; ns, non-significant (p>0.05); using Mann-Whitney test. See also Supplementary Figure 5.

### Niche dynamics are linked with Hoxb5^+^Vwf^+^ HSCs activation

We next sought to characterize the bone marrow niche of Hoxb5^+^Vwf^+^ HSCs. We started by measuring the distance of HSC subsets to blood vessels and MKs (defined as large *Vwf*-GFP^+^ cells). Distances were determined in segmented 3D images from the first timeframe of movies or in tilescans of calvarium BM cavities (**Fig.6A**). As expected, given the high abundance of sinusoidal vessels in BM, both Hoxb5^+^Vwf^+^ and Hoxb5^+^Vwf^-^ cells were equally found in close proximity (<10 µm) to blood vessels, both in homeostasis and 24 hrs post platelet depletion (**Fig.6A-B**). Of note, in homeostasis the Hoxb5^+^Vwf^+^ cells were not preferentially found near MKs when compared with Hoxb5^+^Vwf^-^ cells, in contrast to a previous study^32^. However, 24 hrs post platelet depletion the Hoxb5^+^Vwf^+^ localized closer to MKs (**Fig.6A, C**). While this finding was unexpected given the non-migratory behavior of these cells, we hypothesized that MKs’ distribution post platelet depletion could change instead. We previously reported increased MK numbers in the sternum BM of mice 24 hrs post platelet depletion^29^, and we could confirm this was also happening in the calvarium BM (**Fig.6D**). This is likely explaining the closer proximity Hoxb5^+^Vwf^+^ cells to MKs post platelet depletion. Yet, the fact that Hoxb5^+^Vwf^-^ cells are not found in closer proximity to MKs in the same mice post platelet depletion (**Fig.6B**) suggests the higher proximity of Hoxb5^+^Vwf^+^ cells to MKs is not just stochastically happening due to increased MK numbers and prompted a more detailed analysis of MK behavior. IVM movies from mice post platelet depletion already suggested a highly dynamic megakaryopoiesis, with MKs growing and disintegrating into pro-platelets within the 1 hr observation period (**Fig.4D-E**, green arrows; **Supplementary Movie 4**). Remarkably, the comparison of tilescans from unchallenged mice imaged twice, 24 hrs apart, showed only approximately 30% overlap in MKs, indicating megakaryopoiesis is a highly dynamic process even in homeostasis (**Fig.6E**). Repeated calvarium imaging of chimeric mice stably reconstituted with BM from *Vwf*-TdTomato mice prior to and 24 hrs following platelet depletion confirmed this highly dynamic behavior (**Supplementary** Fig.5B). We previously demonstrated a role of Lepr^+^ perivascular cells in regulating the response of Vwf^+^ HSCs to platelet depletion^29^. Similarly to vasculature, Lepr^+^ cells are highly abundant in BM and therefore, the majority of Hoxb5^+^Vwf^+^ and Hoxb5^+^Vwf^-^ cells were equally found in direct contact with these cells, as observed in thick femur sections from mice in homeostasis (**Fig.6F-G**, **Supplementary** Fig.5C**-D**). All together, these analyses revealed that while Hoxb5^+^Vwf^+^ HSCs essentially do not migrate, the surrounding niche cellular composition may change following acute perturbations in hematopoiesis to regulate the response of these cells to stress.

**Figure 6.**
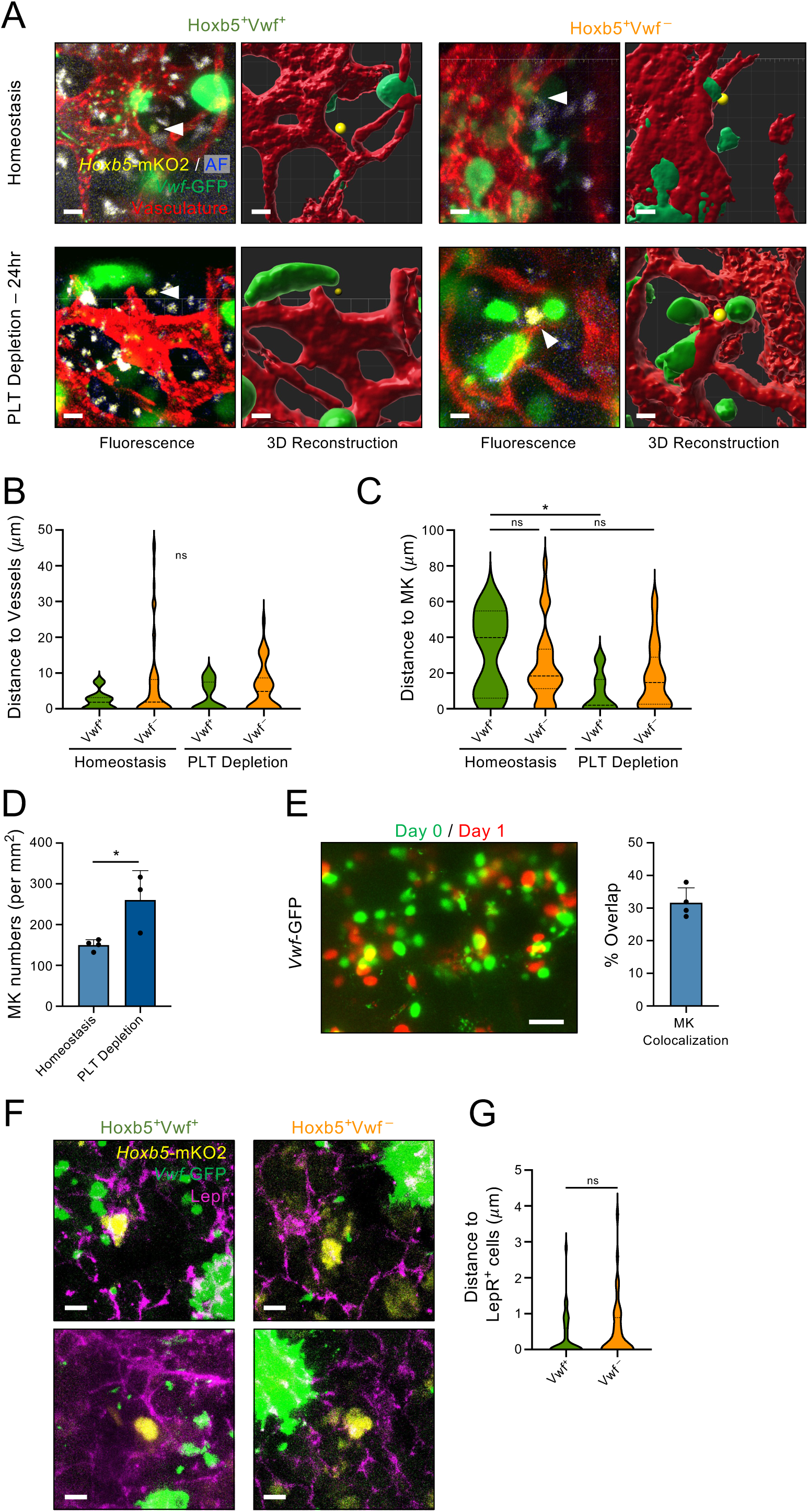
The Hoxb5^+^Vwf^+^ cells locate closer to MKs post platelet depletion due to increased MK numbers and megakarypoiesis. **A-C)** Localization of Hoxb5^+^Vwf^+^ and Hoxb5^+^Vwf^-^ cells in relation to blood vessels and MKs in the calvarium of mice in homeostasis and 24 hrs post platelet depletion. **A)** Representative images and corresponding 3D reconstruction. MKs were identified as large cells expressing *Vwf*-GFP. Blood vessels were labelled by intravenous injection of CD31 and CD144 antibodies. Scale bars represent 10 μm. **B-C)** Distance to vasculature (**B**) and MKs (**C**) for all cells analysed. Data for 7 and 12 Hoxb5^+^Vwf^+^ cells and 68 and 39 Hoxb5^+^Vwf^-^ cells from 12 mice in homeostasis and 11 mice analysed 24 post platelet depletion, respectively. *, p<0.05; ns, non-significant (p>0.05); using Kruskal-Wallis test with Dunn’s multiple comparisons. **D)** Number of MKs in the calvarium BM of mice in homeostasis or 24 hrs post platelet depletion. N=4 and 3 mice in homeostasis and post platelet depletion, respectively. *, p<0.05 using t-test. **E)** Serial imaging of MKs in the calvarium BM in homeostasis with 24 hrs interval showing high megakaryopoiesis dynamics in homeostasis and post platelet depletion. Left panel shows representative overlayed images of the same mouse BM at both time points. Images from same mouse taken at different time points were aligned based on vasculature and bone autofluorescence signal. Right panel shows the percentage of overlap in MK localization between images taken at both time points. N=4. Scale bar represents 100 μm. **F-G)** Localization Hoxb5^+^Vwf^+^ and Hoxb5^+^Vwf^-^ cells in relation LepR^+^ perivascular cells in the femur of mice in homeostasis. **F**) representative images and **G**) and quantification of the distance to LepR^+^ cells, measured in 3D tile scans of the bone marrow. Data from 35 Hoxb5^+^Vwf^+^ cells and 41 Hoxb5^+^Vwf^-^ cells from 3 mice analysed. Scale bars represent 5 μm. See also Supplementary Figure 5.

### Hoxb5^+^Vwf^+^ HSCs remain non-migratory during ageing

Ageing is associated with the expansion of the HSC compartment, increased HSC bias towards the platelet/MK lineage^39^, and increased platelet numbers in mice (**Supplementary** Fig.6A). Unlike acute platelet depletion, changes in hematopoiesis develop slowly and continuously, and we wondered whether it would be linked to changes in Hoxb5^+^Vwf^+^ HSCs’ interactions with their niche. To assess this, we investigated the behavior and localization of Vwf^+^ HSCs in 1 year old mice, when the main hallmarks of ageing in HSCs and hematopoiesis are already observed^40^. The analysis of the HSC compartment of ageing *Hoxb*5-mKO2^Tg/+^*Vwf*-GFP^Tg/+^ mice revealed a 15-fold expansion in the number of total SLAM-HSCs, particularly affecting the Hoxb5^+^Vwf^+^ compartment, which became the most represented HSC subset, comprising 66.4±6.6% of all SLAM-HSCs (**Fig.7A and** **Supplementary** Fig.6B**-C**). Furthermore, in agreement with previous studies^41^, both Hoxb5^+^Vwf^+^ and Hoxb5^+^Vwf^-^ SLAM-HSC subsets from ageing mice showed increased EPCR expression, in comparison to young mice, with the increase more prominent in the Hoxb5^+^Vwf^-^ subset (**Supplementary** Fig.6D**-E**). Of note, analysis of cell populations solely defined by the expression of *Hoxb5* and *Vwf* reporters revealed a small but significant increase in the frequency of phenotypic HSCs (**Fig.7B and** **Supplementary** Fig.6F), which constituted 87.3±3.9% and 53±8.9% of the total Hoxb5^+^Vwf^+^ and Hoxb5^+^Vwf^-^ cells, respectively. No major differences were observed in the cellular composition of the remaining non-HSC cells within the total Hoxb5^+^Vwf^+^ and Hoxb5^+^Vwf^-^ populations, despite a small increase in the proportion of Hoxb5^+^Vwf^+^ MkPs (**Supplementary** Fig.6G), overall confirming the suitability of our double reporter strategy to identify phenotypic Vwf^+^ HSC also in ageing mice.

**Figure 7.**
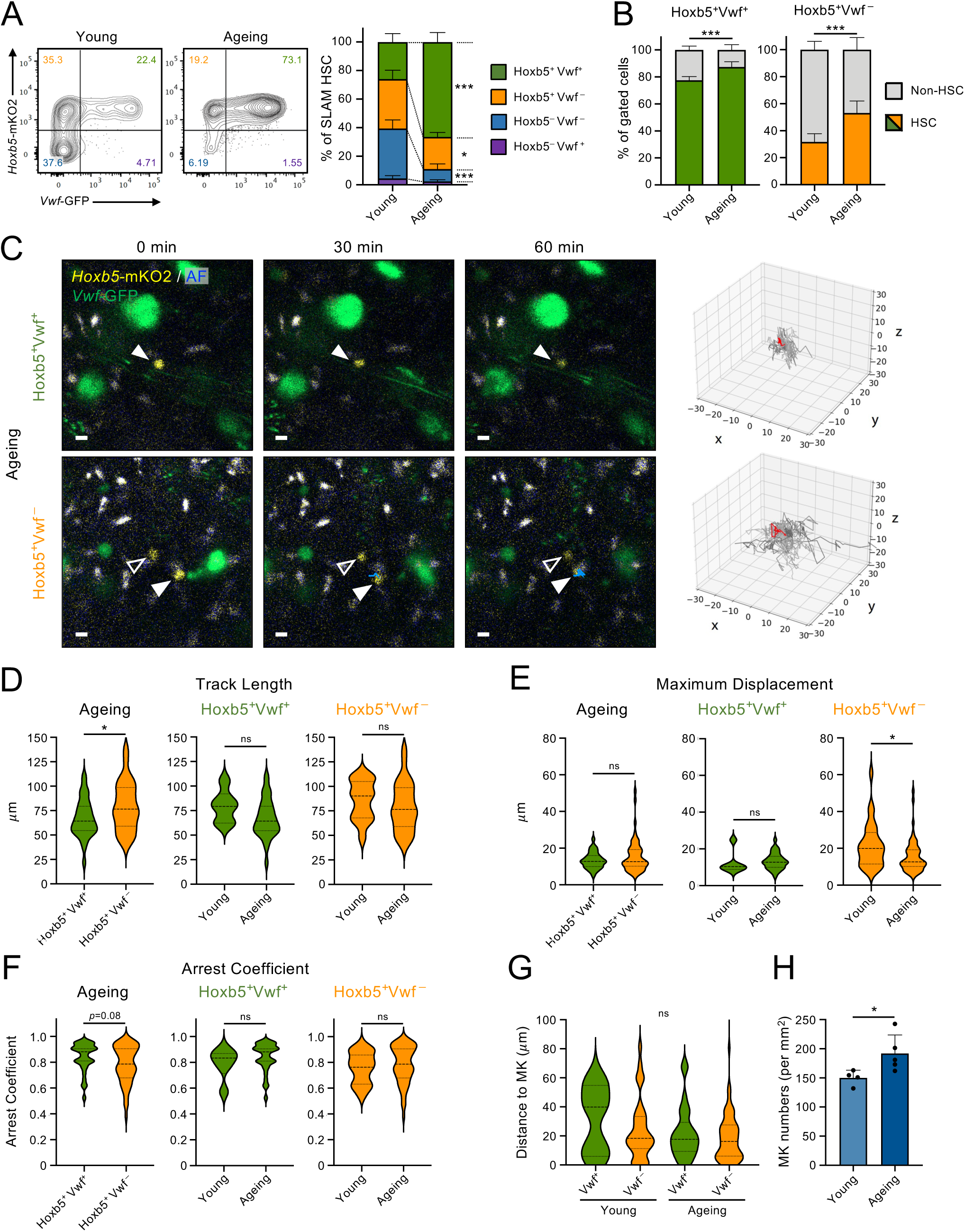
The Hoxb5^+^Vwf^+^ cells retain a highly immobile behaviour during ageing. **A)** Flow cytometric analysis of the *Hoxb5*-mKO2 and *Vwf*-GFP reporters’ expression in phenotypically defined SLAM-HSCs from young and ageing mice. Representative FACS profiles of SLAM-HSC subsets defined by *Hoxb5*-mKO2 and *Vwf*-GFP expression (left) and mean±SD frequencies from 6 young and 9 ageing mice in 4 independent experiments (right). Numbers in quadrants indicate the frequency of the gated subset within the SLAM-HSC population. **(B)** mean±SD frequency of SLAM-HSCs within the Hoxb5^+^Vwf^+^ and Hoxb5^+^Vwf^-^ populations defined exclusively based on the expression of the reporter transgenes. Data from 6 young and 9 ageing mice from 4 independent experiments. **(C-F)** The calvarium BM of double reporter 1 year old mice was imaged by intravital microscopy (IVM) for 1 hr. **C)** Selected timeframes at the indicated time points, from 3D time-lapse IVM, showing the dynamics of representative Hoxb5^+^Vwf^+^ and Hoxb5^+^Vwf^-^ cells (indicated by filled and empty arrowheads). The trajectory of the cells during the imaging period is indicated in blue. Scale bars represent 10 μm. Rose plots on the right show the 3-dimensional tracks (in μm) of all cells analysed (grey), with the cells depicted in the left images by a filled arrowheads highlighted in red. Tracks for each cell were normalized for position of origin. **(D-F)** Track parameters extracted from the 3D time-lapse datasets of single Hoxb5^+^Vwf^+^ and Hoxb5^+^Vwf^-^ cells (in **C**). Violin plots depict track length **(D)**, maximum displacement **(E)**, and arrest coefficient **(F)**. Data in **(D-F)** is from 6 and 32 Hoxb5^+^Vwf^+^ cells, and 28 and 56 Hoxb5^+^Vwf^-^ cells from, 12 mice in homeostasis and 5 ageing mice, respectively. **G)** Distance of Hoxb5^+^Vwf^+^ and Hoxb5^+^Vwf^-^ cells to MKs, in young and 1 year old mice. Data from 7 and 43 Hoxb5^+^Vwf^+^ and 68 and 178 Hoxb5^+^Vwf^-^ cells, from 12 young mice and 5 ageing mice, respectively. **H)** Number of MKs in the calvarium BM of young (N=4) and 1 year old mice (N=5). *, p<0.05; ***, p<0.001; ns, non-significant (p>0.05); using 2-way ANOVA with Sydak’s multiple comparisons **(A)**, t-test **(B, H)**, Mann-Whitney test **(D-F)** or Kruskal-Wallis test with Dunn’s multiple comparisons **(G)**. Data depicted for young mice are the same shown in Fig.5-6. See also Supplementary Figures 6 and 7.

We next compared the dynamic behavior of Hoxb5^+^Vwf^+^ and Hoxb5^+^Vwf^-^ in the calvarium BM of ageing mice using the same experimental approach previously applied to young mice (**Fig.7C; Supplementary Movies 7-8**). Remarkably, and similarly to our observations in young mice, ageing Hoxb5^+^Vwf^+^ cells exhibited a stationary yet wiggly behavior, characterized by oscillatory movements that lead to mean track lengths of 75µm/hr, but without resulting in a significant displacement of the cells within the BM niche. Consistently, these cells displayed a high arrest coefficient and low linearity coefficient (**Fig.7D-F and** **Supplementary** Fig.7A**-D**). While in ageing mice the Hoxb5^+^Vwf^-^ population still has a more heterogenous cellular composition when compared with the Hoxb5^+^Vwf^+^ population (**Fig.7B and** **Supplementary** Fig.6F**-G**) aged Hoxb5^+^Vwf^-^ were less motile than their young counterparts, indicated by their reduced maximum displacement and linearity coefficient (**Fig.7E and** **Supplementary** Fig.7C**-D**). Similarly to what we observed in young mice, ageing Hoxb5^+^Vwf^+^ cells do not preferentially localize in close proximity to MKs, when compared with Hoxb5^+^Vwf^-^ cells (**Fig.7G**), despite a small but significant increase in MK numbers in the bone marrow of aged mice (**Fig.7H**) and the increased numbers of Vwf^+^ HSCs (**Fig.7A and** **Supplementary** Fig.5B**-C**). This further supports that the closer proximity of Hoxb5^+^Vwf^+^ HSCs to MK post platelet depletion is stress specific and not only a stochastic phenomenon resulting from increased MK numbers.

## DISCUSSION

Here we described a new strategy to identify highly enriched endogenous HSCs in mouse BM, based exclusively on co-expression of the *Hoxb5*-mKO2 and *Vwf*-GFP reporters. Combined reporters’ expression identifies a rare population in whole BM constituted of 83% SLAM-LSKs, to our knowledge the highest frequency of phenotypic HSCs so far reported when using only transgenic reporter alleles^9,10,12-16^. The high purity of LT-HSCs among the Hoxb5^+^Vwf^+^ cells was further demonstrated functionally with highly efficient reconstitution of primary and secondary recipient mice, and at transcriptional level as evidenced by their HSC and repopulation scores. In contrast, Hoxb5^+^Vwf^-^ cells represent a more heterogenous population that in addition to LT-HSCs also includes a significant proportion of MPP1/ST-HSCs and MPP4 cells. In line with the high frequency of MPP1/ST-HSCs within the Hoxb5^+^Vwf^-^ compartment, these cells efficiently repopulate primary recipient mice but are less efficient in reconstituting their BM HSC compartment, and in subsequent reconstitution of secondary recipients.

Similarly to other reporter strategies, Hoxb5^+^Vwf^+^ cells only represent a fraction of LT-HSCs. Direct comparisons between HSC-reporter mice are currently impossible due to fluorescence overlap of the reporter genes used. Therefore, while the identification of cells based on transcript levels may partially differ from reporter-transgene expression, particularly when reporter activation depends on Cre-mediated recombination events, our scRNA-Seq analysis approach provides the current best strategy to compare HSC definitions used to identify HSCs *in situ*. This analysis revealed that while >95% of Hoxb5^+^Vwf^+^ cells are included in the populations defined by other reporter-based strategies, they only constitute a small fraction of these populations. Nevertheless, the population defined by *Hoxb5*-mKO2 and *Vwf*-GFP expression comprises the totality of the Vwf^+^ LT-HSCs and therefore represents a functionally relevant subset of HSCs that are highly potent and have a platelet differentiation bias^24,26^. Importantly, we found little overlap in the cells identified between each of the other HSC definitions tested and did not find any cells Identified by all definitions together, highlighting significant differences between HSC reporter strategies. Furthermore, we observed different spreading of the identified cells across the HSPC landscape, indicating different types and proportions of non-HSCs are also being identified, affecting the comparisons of results obtained with these reporter mice.

Previous studies reported conflicting evidence regarding the behavior of HSCs *in vivo*, both in homeostasis and following chemotherapy and mobilization treatments^10,16^. Our time-lapse imaging of Hoxb5^+^Vwf^+^ cells in the calvarium BM revealed their non-migratory behavior, which despites the low level of overlap between transcriptional definitions is however in line with the low motility previously described for MFG^10^ and *Hlf*-tdTom^hi^ cells^15^ in homeostasis. Although the cells are not migratory, they show a wiggling, oscillatory movement that does not result in net-displacement. This wiggling behavior was previously described but only in transplantation settings, and suggests an active engagement of these cells with the niche^20^. In contrast, Hoxb5^+^Vwf^-^ cells show a more heterogenous behavior including both immobile and migratory cells, which likely reflects the heterogeneous composition of the Hoxb5^+^Vwf^-^ population, including a higher proportion of downstream progenitors. This suggests that immature progenitors are more motile when compared to LT-HSCs, which is consistent with the more static behavior of MFG cells when compared with the broader *Mds1*-GFP^+^ population, that is highly represented by progenitor cells^10^, and with earlier studies comparing the dynamics of human stem and progenitor cells following transplantation in non-irradiated recipients^18^. Contrarily to our observations, Upadhaya et al., previously reported higher motility of cells identified by *Pdzk1ip1*-CreER mediated recombination of a R26^LSL-Tom^ reporter (Pdzk1ip1-Tom^+^ cells). In this study, only 13% of Pdzk1ip1-Tom^+^ cells show a static behavior and the majority are migratory and characterized by an intermittent behavior, with phases of progressive motion separated by occasional short periods of confined random motion^16^. Although our imaging window was shorter this is not likely to explain the non-migratory behavior of Hoxb5^+^Vwf^+^ cells since the observed sporadic phases of confined random motion of *Pdzk1ip1*^+^ cells^16^ are not compatible with the 1 hr periods of arrested behavior we observed here. The different behavior of Hoxb5^+^Vwf^+^ and Pdzk1ip1-Tom^+^ cells may therefore reflect different populations of cells being analyzed or result from the requirement for Tamoxifen administration to induce reporter activation in this model, which may interfere with HSCs dynamics and lead to variable levels of HSC purity within the labeled cell population^42^. We can however not exclude that the *Hoxb5*^+^*Vwf*^+^ cells might constitute the 13% *Pdzk1ip1*^+^ cells showing a static behavior^16^.

Moreover, we provided here the first analysis to our knowledge of HSC behavior during ageing. The analysis of 1 year old mice, when the HSC compartment had already significantly expanded, and hematopoiesis shows the main phenotypes associated with ageing^40^, similarly showed Hoxb5^+^Vwf^+^ cells maintained their non-migratory behavior. Previously, *Hlf*-tdTom^hi^ cells were shown to be migratory in the neonatal period and subsequently become stationary in the adult BM^15^. Together those and our data suggest that the non-migratory behavior is instated at the transition between fetal and adult hematopoiesis and persists throughout life.

Our new HSC reporter strategy also showed high stability in stress hematopoiesis enabling the longitudinal analysis of HSC behavior in homeostasis and following perturbations to steady state. Remarkably, 24 hrs post platelet depletion, when over 60% of Vwf^+^ SLAM-HSCs are actively proliferating^29^, the Hoxb5^+^Vwf^+^ cells retain their static behavior. Mobilization or 5-FU treatments were previously shown to induce the motility of MFG cells^10^, potentially due to stronger effects on hematopoiesis but also on the BM niche. Nevertheless, despite the overall increased motility the response of MFG cells to these stresses was heterogeneous with a fraction of the cells displaying limited displacement, compatible with the partial overlap between Hoxb5^+^Vwf^+^ and MFG cells. The higher LT-HSC enrichment observed within the Hoxb5^+^Vwf^+^ population together with the higher functionality of Vwf^+^ HSCs^24,26^ suggests that the immobile behavior in homeostasis and stress might be a property of the most potent and immature HSCs. Surprisingly, Pdzk1ip1-Tom^+^ cells, that are already more dynamic in homeostasis, become less motile post mobilization^16^. Whether the highly mobile cells observed in other studies represent different HSC subsets, downstream progenitors included within the identified populations or, stress-specific response behaviors will require further investigation.

Although Vwf^+^ HSCs have been previously suggested to localize nearer MKs^32^, we did not find a preferential localization of Hoxb5^+^Vwf^+^ HSCs closer to those cells, compared to Hoxb5^+^Vwf^-^ cells, which would have been expected. However, despite their immobile behavior, Hoxb5^+^Vwf^+^ cells appear closer to MKs after platelet depletion, partially due to increased megakaryopoiesis and MK numbers in BM. The increased proximity post platelet depletion is not entirely stochastic as it was only observed for Hoxb5^+^Vwf^+^ cells. Understanding why Hoxb5^+^Vwf^+^ but not Hoxb5^+^Vwf^-^ cells appear closer to MKs will require further investigation and the recent studies describing an alternative pathway for fast MK differentiation during regeneration and ageing^28,43-45^ raise the possibility that Hoxb5^+^Vwf^+^ HSCs may quickly and directly generate MKs post platelet depletion thereby explaining the close localization of these cells. In fact, sequential imaging of the calvarium BM revealed notorious changes in MK localization uncovering a new aspect of niche dynamics in response to stress. MKs play a critical role in instructing HSC quiescence through TGFb and PF4^30,31,46^. Following cell cycle activation in response to platelet depletion Vwf^+^ HSCs rapidly return to quiescence and the closer proximity of Hoxb5^+^Vwf^+^ HSCs to MKs at 24 hrs may underly a role of MKs themselves in reestablishing HSC quiescence. Both Hoxb5^+^Vwf^+^ and Hoxb5^+^Vwf^-^ cells were found adjacent to blood vessels and LepR^+^ perivascular cells, in line with the high abundance of these niche cells^4^ and reflecting the need to identify markers to explore niche cells heterogeneity to precisely define HSC niches. Previous studies suggested the existence of different niches supporting quiescent and activated HSCs^4,8,10,33,34^. However, despite the high proliferation rate after platelet depletion Hoxb5^+^Vwf^+^ HSCs retain a highly immobile behavior in the BM niche. Although we cannot exclude that in different scenarios of stress hematopoiesis HSCs may relocate to a different niche, our findings unlink cell cycle activation and the requirement to occupy an alternative niche eliciting proliferation. Nevertheless, the surrounding niche may change both molecularly and in cellular composition as observed here with the expansion of MKs, indicating the need for a better understanding of niche remodeling in situations of perturbed hematopoiesis.

We reported a new strategy that stably identifies highly enriched LT-HSC *in situ* and *in vivo*. While the increased enrichment observed here leads to the identification of only a fraction of HSCs and to extremely limited numbers of cells that can be found *in situ*, the independence of Cre recombination will allow this reporter to be combined with other transgenic systems to model hematological diseases and investigate HSC-niche interactions in these scenarios.

## METHODS

### Mice

All mice were bred and maintained in accordance with the UK Home Office regulations. All procedures were performed under project licenses PP9504146 and PP6427565 approved by the Imperial College London Animal Welfare and Ethics Review Body. *Hoxb5*-mKO2^12^, *Vwf*-GFP^26^ and *Vwf*-TdTomato^24^ mice have been previously described. B6.SJL-*Ptprc*^a^*Pepc*^b^/BoyJ (CD45.1) mice (The Jackson Laboratory, 002014) were used as transplantation recipients. *Hoxb5*-mKO2 mice were generated and kindly provided by Dr Dónal O’Carroll (University of Edinburgh) and Dr Kamil Kranc (ICR). *Vwf*-GFP mice were kindly provided by Prof Sten Eirik Jacobsen and Prof Claus Nerlov (University of Oxford). Bone marrow from *Vwf*-TdTomato mice was kindly provided by Dr Simon Mendez-Ferrer (University of Cambridge). All mouse lines were backcrossed for at least 6 generations onto a C57BL/6 genetic background and littermate controls were used in all experiments. Young mice were used at 7-14 weeks of age, and ageing mice were used at 59-70 weeks of age. Mice of both sexes were used in all experiments.

### Platelet depletion

GPIbα-antibody mediated platelet depletion was performed as previously described^29^. Briefly, platelet depletion was induced by one intra-venous (IV) administration of the anti-GPIbα antibody (R300; Emfret Analytics) at 2μg/g body weight at the indicated time points before analysis.

### Competitive bone marrow transplantation

*In vivo* limiting dilution analysis was performed by transplanting 5, 10 and 15 Hoxb5^+^Vwf^+^ or Hoxb5^+^Vwf^-^ cells (CD45.2) along with 2.5x10^5^ unfractionated support/competitor CD45.1 bone marrow cells into lethally irradiated (10 Gy, split dosage of 5 Gy each) recipient mice (CD45.1) of 8-12 weeks of age. Live (DAPI^-^) Hoxb5^+^Vwf^+^ or Hoxb5^+^Vwf^-^ cells were isolated by flow cytometry, based solely on reporters’ expression and without any other HSC markers. Flow cytometric analysis of CD45.1 and CD45.2 contribution to mature peripheral blood lineages was performed at 4, 8, 12 and 16 weeks after transplantation. Mice were considered reconstituted if showing a CD45.2 contribution to platelet, myeloid and lymphoid lineages of at least 0.1%. Donor contribution for the platelet/megakaryocyte lineage in peripheral blood was performed based on expression of the *Vwf*-GFP transgene, as previously described^24,26^. At 16 weeks post transplantation CD45.2 repopulation of the HSC compartment in bone marrow was investigated and 2x10^6^ unfractionated bone marrow cells from the mice initially transplanted with 10 cells was re-transplanted into secondary recipients to evaluate long-term reconstitution capacity in peripheral blood and bone marrow at 16 weeks post transplantation.

### Flow Cytometry

For bone marrow analysis, the femurs, tibia, hip bones and sternum were crushed to prepare single cell suspensions. For peripheral blood analysis, blood was collected in EDTA coated tubes (Sarstedt) and centrifuged at 100g, for 10 min at room temperature (RT) to isolate the platelets fraction. The remaining blood was then mixed with Dextran (1%; Sigma) and incubated for 40 min at 37°C to separate red blood cells. The top fraction (containing the mononuclear cells was further incubated with NH_4_Cl solution (Stem Cell Technologies) for 2 min at RT to lyse the remaining red blood cells. Mouse BM single cell suspensions and peripheral blood cells were Fc-blocked and stained with titrated antibodies for mouse antigens (**Supplementary Table 1**). 7-Amino-Actinomycin D (7AAD; Sigma), 4′,6-diamidino-2-phenylindole (DAPI; Invitrogen) or Fixable Viability Stain 510 (BD Biosciences) were used for dead cells exclusion. Compensations were initially performed with BD CompBeads (BD Biosciences) followed by manual adjustments using fluorescence-minus-one (FMO) controls for all channels. FMO controls and negative populations were used to set gates. Absolute cell numbers were defined as cells per 2 legs (each including femur, tibia and hip bone). FACS analyses were performed on a BD Fortessa X20 (BD Biosciences) and subsequently analysed with FlowJo Software (BD Biosciences). Cell sorting experiments were performed on a BD FACS Aria III (BD Biosciences), with a mean cell purity of 98.2%.

### Intravital microscopy

Intravital imaging was performed as previously described^17^. Briefly, mice were imaged using a water-immersion 20X objective on a Zeiss LSM980 Axio Examiner.Z1 upright confocal microscope equipped with a 405nm, 488nm, 561 nm and 639 nm diode lasers, a 594 nm diode pumped solid state (DPSS) laser, an internal spectral detector array and 6 external detectors (2 non-descanned detectors (NDD), 2 Gallium Arsenide Phosphide Photomultiplier Tubes (GaAsP PMT) and 2 nosepiece GaAsP PMT). For the detection of GFP and mKO2 signals, excitation was performed using the 488nm and 561nm lasers, respectively, and acquired in 2 separate tracks (1 track for GFP and AF647 and the second track for mKO2 and autofluorescence). Autofluorescence signal was detected on the AF594 channel (with 561 nm excitation only) and used to identify real mKO2 signal. Fluorescently labelled antibodies against Pecam1/CD31 and VE-Cadherin/CD144 (**Supplementary Table 2**) were injected intravenously 15-30 min prior to imaging, to detect vasculature (**Supplementary table 2**). In some experiments vasculature was detected by intravenous injection of 650µg of Cy5-labelled 500KDa Dextran (Nanocs) per mouse. Time-lapse imaging was performed by acquiring 3D images every 3 minutes. Due to bleaching of the mKO2 reporter and to allow serial imaging of the cells at different time points most cells were not tracked for more than 1 hr and therefore all movies were capped at 1 hr. Serial imaging of *Vwf*-GFP and *Vwf*-tdTomato mice calvarium to observe MKs at different time points confirmed the dynamic megakaryopoiesis observation does not result from possible artifacts due to re-imaging GFP^+^ cells by IVM.

### Intravital Imaging analysis

Hoxb5^+^Vwf^-^ and Hoxb5^+^Vwf^+^ cells were identified in Imaris (v9.7.2, Oxford Instruments) using the manual tracking tool on the KuO channel in both movies and tilescans, and cell positions were exported and plotted using matplot3d in Jupyter Notebook. X, Y and Z drift from one frame to the next was corrected in movies using the Fast4Dreg macro^47^. Track parameters and formulas used were previously described^48^. Hoxb5^+^Vwf^-^ and Hoxb5^+^Vwf^+^ cells were segmented using the Imaris “spots” function. MKs and vessels were also segmented in Imaris using the “surface detection” function, an absolute intensity threshold that varied per image and a voxel volume threshold. For tilescans analysis, the boundaries of the bone were defined using the autofluorescence channel for every Z-plane, and the bone was then cropped in every channel using a custom Fiji macro. MKs were defined as GFP+ cells with a volume >980 µm^2^ (equivalent to a diameter >12.3 µm). MK quantification and overlap in calvarium tiles cans were analysed with ImageJ/Fiji software.

### Imaging of thick bone marrow sections

Undecalcified femurs were fixed in 4% paraformaldehyde at 4°C, overnight and with rotation. Bones were then washed in incubated in a sucrose gradient of 10% (1 hr) 20% (1 hr) and 30% (overnight), before embedding and cryopreserved in OCT compound (Leica). Bones were sectioned on a Leica CM3050S cryostat equipped with the CryoJane Tape transfer System (Leica). 50 μm thick sections were affixed on slides pre-coated with Norland Optical Adhesive 63 (Norland Products) and exposed to 2 flashes of UV light. Tissue sections were blocked with 20% normal donkey serum for 1 hour and stained with primary antibodies for 2 days at 4°C, then stained with secondary antibodies (**Supplementary Table 2**) for 3 hours at 4°C, mounted in Prolong Glass Antifade mountant (ThermoFisher), cured for 24 hours at RT in the dark and imaged on a Leica Stellaris 8 inverted microscope using a 20X objective. HoxB5 signal was detected using the AF555 channel using Leica TauSense Imaging Tool and TauGating function to separate the true signal (0.2 – 1ns signal lifetime gate) from background autofluorescence (1 ns onwards). Autofluorescence was further detected on an empty 405 channel. Z-stacks with 2μM intervals were acquired to capture the full depth of the tissue section. Images were analysed on ImageJ/Fiji software.

### RNA sequencing analysis

Identification of *Hoxb5*^+^*Vwf*^+^ cells in Nestorowa *et al* dataset: The processed data was downloaded from the Nestorowa dataset^35^ link (https://blood.stemcells.cam.ac.uk/single_cell_atlas.html), which contains 8 classified stem and progenitor cell populations, 1644 cells, and force-directed graph. Stem and progenitor cell populations were classified based on single cell index sorting data and defined as: LT-HSC (Lin^-^Sca1^+^cKit^+^Flt3^-^CD34^-^), ST-HSC (Lin^-^Sca1^+^cKit^+^Flt3^-^CD34^+^CD48^-^ CD150^-^), MPP1 (Lin^-^Sca1^+^cKit^+^Flt3^-^CD34^+^CD48^-^CD150^+^), MPP2 (Lin^-^Sca1^+^cKit^+^Flt3^-^ CD34^+^CD48^+^CD150^+^), MPP3 (Lin^-^Sca1^+^cKit^+^Flt3^-^CD34^+^CD48^+^CD150^-^), LMPP (Lin^-^ Sca1^+^cKit^+^Flt3^+^CD34^+^), MEP (Lin^-^Sca1^+^cKit^+^CD16/32^-^CD34^-^), CMP (Lin^-^Sca1^+^cKit^+^CD16/32^-^CD34^+^) and GMP (Lin^-^Sca1^+^cKit^+^CD16/32^+^CD34^+^). First, the expression of two genes, *Hoxb5* and *Vwf*, were extracted from the log-normalized expression matrix. Based on the relative expression values of these two genes in all 8 progenitor cell populations, hard thresholds of 3 and 3 were chosen respectively to define *Hoxb5*^+^*Vwf*^+^ cells. The defined *Hoxb5*^+^*Vwf*^+^ and *Hoxb5*^+^*Vwf*^-^ cells were then displayed in the landscape using the scanpy.tl.draw_graph^49^ function to examine the overlap with LT-HSCs.

HSC and repopulation scores: The HSC Score is calculated based on the expression values of the 103 molecular overlapping population (MolO) signature genes using the hscScore model^36,37^. The *Vwf* gene itself was excluded from the hscScore calculations because it is also being used to define the cells. The Repopulation signature score was calculated based on the expression values of 23 relevant Repopulation signature genes^38^ using the scanpy.tl.score_genes function. Wilcoxon test was used to calculate the significance of differences. Asterisks denote a significant difference (*** p ≤ 0.001; ** p ≤ 0.01; ns p > 0.05). All violin plots were performed using ggplot2 package (https://ggplot2.tidyverse.org) in R.

Comparison between HSCs definitions: Four additional HSC definitions were introduced, which included *Pdzk1ip1*^+^, *Tcf15*^+^, *Ctnnal1*^+^*Hlf*^+^ and *Mds1*^+^*Flt3*^-^. The expression of *Pdzk1ip1*, *Tcf15*, *Ctnnal1, Hlf* and *Flt3* was obtained directly from the downloaded processed data. Thresholds defining each positive/negative cell type were determined based on their expression in eight different progenitor cell populations (6, 3, 4 5 and, 3 respectively). Since the downloaded processed data only contains the expression of the *Mecom* complex locus and not of *Mds1* or *Evi1* genes within this locus, re-alignment was performed to retrieve the expression of *Mds1*. The coordinates of the *Mds1* exons were extracted from the mouse Ensemble genome browser (GRCm38) based on the description of the mouse reporter transgene^10^ Only the first two exons of *Mds1,* common to both transcripts produced from the *Mds1* promoter were used to more accurately determine *Mds1* expression. A threshold of 2 was used to identify *Mds*1^+^ cells. This was also compared with the expression of the whole *Mecom* locus. Raw sequencing data were downloaded from GEO according to the GEO accession number (GSE81682). To make the results consistent with the downloaded processed dataset, it was then realigned to the GRCm38 reference genome with 92 spike-ins developed by the External RNA Control Consortium (ERCC) using STAR^50^ (version 2.7.3a). Count matrix was generated using featureCounts^51^ (version 2.0.8), focusing only on Mds1 and Evi1. Venn plots were generated using the R package venn (https://github.com/dusadrian/venn) were used to examine the number of intersecting cells between the 5 HSCs definitions. The percentage of intersections was calculated by dividing the number of intersecting cells by the total number of cells in the defined category, which was then displayed in a heat map **Analysis of Blood cell parameters.** Mouse blood parameters were analysed using a XP-300 (Sysmex) automated blood cell analysers.

### Statistical analysis

Statistical significance was determined by two-sided t-test, 1- or 2-way ANOVA combined with Tukey’s or Sidak’s multiple comparisons tests, Mann-Whitney test and Kruskal-Wallis test with Dunn’s multiple comparisons test, as indicated in figure legends. Statistical tests were performed with GraphPad Prism software. Statistical analysis of LDA *in vivo* experiments was performed with the ELDA software. Experiments were not randomized. The investigators were not blinded to mouse/sample allocation during experiments and outcome assessment.

## ACKNOWLEDGEMENTS

We thank S.E. Jacobsen (University of Oxford & Karolinska Institute) and C. Nerlov (University of Oxford) for providing the *Vwf*-GFP mice; S. Méndez-Ferrer (University of Cambridge) for providing *Vwf*-tdTomato BM; J.E. Rowling and L. Zarate Garcia from The Imperial College FACS Facility; M.Tisi from Imperial College FILM facility and the Imperial College Central Biomedical Services. This work was supported by a Wellcome Trust PhD Fellowship to M.A.S. (222304/Z/21/Z); Wellcome Trust and The Royal Society Sir Henry Dale Fellowship to T.C.L. (210424/Z/18/Z); Kay Kendall Leukaemia Fund Project Grant to T.C.L. (KKL1379); Wellcome Trust Investigator award (212304/Z/18/Z) to C.L.C.; Cancer Research UK Programme Foundation Award (C36195/A26770) to C.L.C.; a Royal Society International Exchange Grant (IEC\R1\180061) to C.L.C. and a Fondation Alcea grant to C.L.C. Wellcome Trust Accelerator Award (313665/Z/24/Z) to A.J. RNA Sequencing analysis was conducted in the Göttgens lab, supported by by Wellcome Trust (206328/Z/17/Z), MRC (MR/W031663/1), Wellcome Trust [203151/Z/16/Z, 203151/A/16/Z] and the UKRI Medical Research Council [MC_PC_17230]. For the purpose of open access, the author has applied a CC BY public copyright licence to any Author Accepted Manuscript version arising from this submission. K.R.K. is a Cancer Research UK (CRUK) Programme grant holder. The Kranc laboratory is funded by CRUK, the Barts Charity, Blood Cancer UK, Medical Research Council (MRC), and the Institute of Cancer Research.

## AUTHORSHIP

Conceptualization: T.C.L. and C.L.C.; Validation: M.A.S, C.L.C. and T.C.L.; Formal analysis: M.A.S, Q.L., C.M., C.L.C. and T.C.L.; Investigation: M.A.S, Q.L., S.G.A., C.M., A.J., J.X., N.W., C.L.C. and T.C.L.; Resources: D.O.C. and K.K. for providing the *Hoxb5*-mKO2 mice; Data curation: M.A.S., Q.L., C.L.C. and T.C.L.; Writing original manuscript: M.A.S, C.L.C. and T.C.L.; Visualization M.A.S, Q.L., C.L.C. and T.C.L.; Supervision and input on experimental design and analysis: T.C.L., C.L.C., B.G., K.K., N.W.; Project Administration: T.C.L. and C.L.C.; Funding Acquisition: T.C.L. and C.L.C. All authors reviewed and edited the final manuscript.

---

**Supplementary Figure 1.**
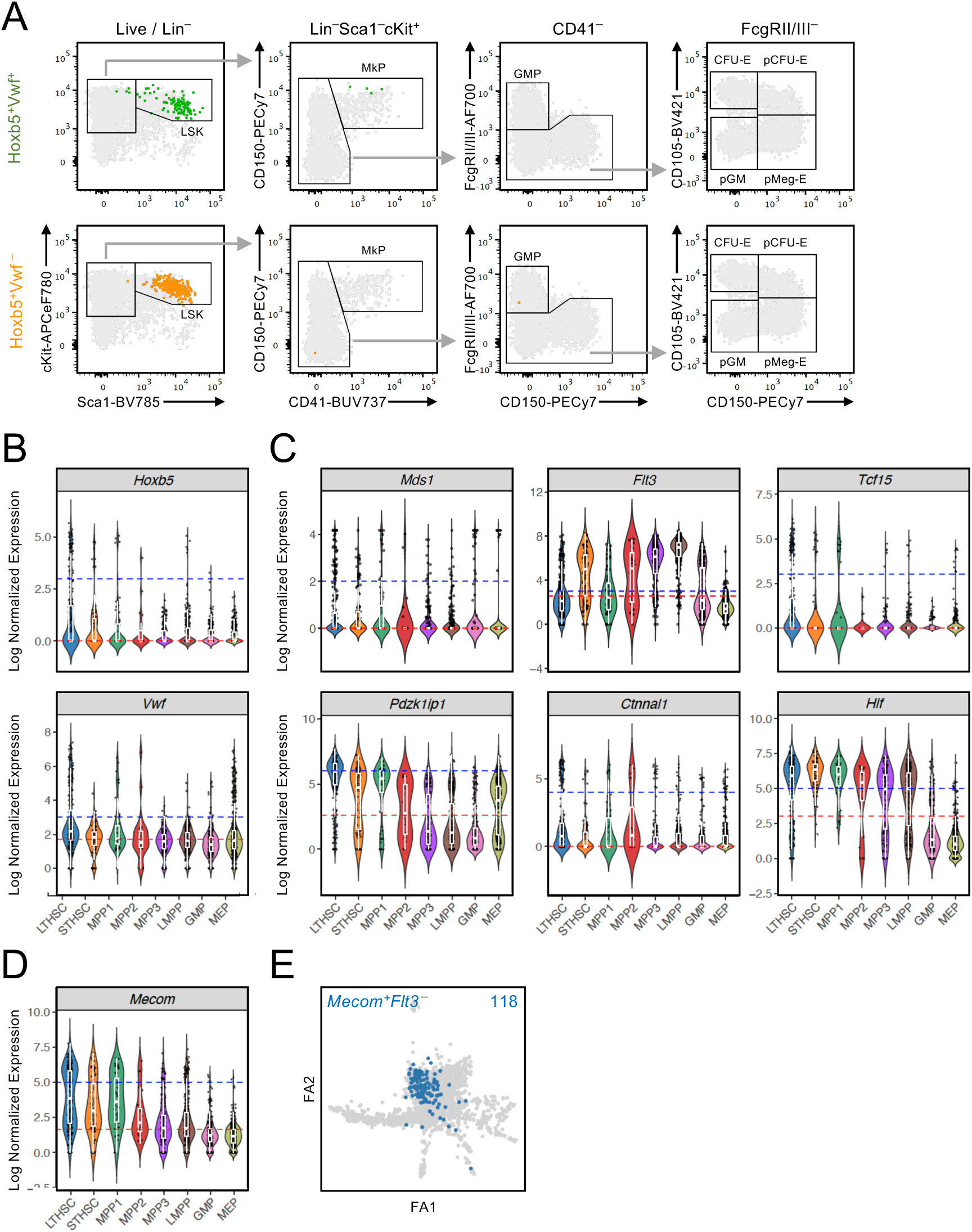
Identification of cell populations by flow cytometry and single cell RNA sequencing. Related to main Figures 1-2. **A)** Representative FACS profiles and gating strategy of myeloid progenitors in BM. **B-D)** Normalized expression of the indicated genes within all populations identified based on immunophenotype from Index sorting, used to define the thresholds for the identification of positive and negative cells for those specific genes. Red dashed line represents the median gene expression across the whole dataset. Blue dashed line represents the threshold used for the identification of cells. **E)** Projection of *Mecom*^+^*Flt3*^-^ cells on the Nestorowa LSK landscape. Number in graph indicates the number of cells identified.

**Supplementary Figure 2.**
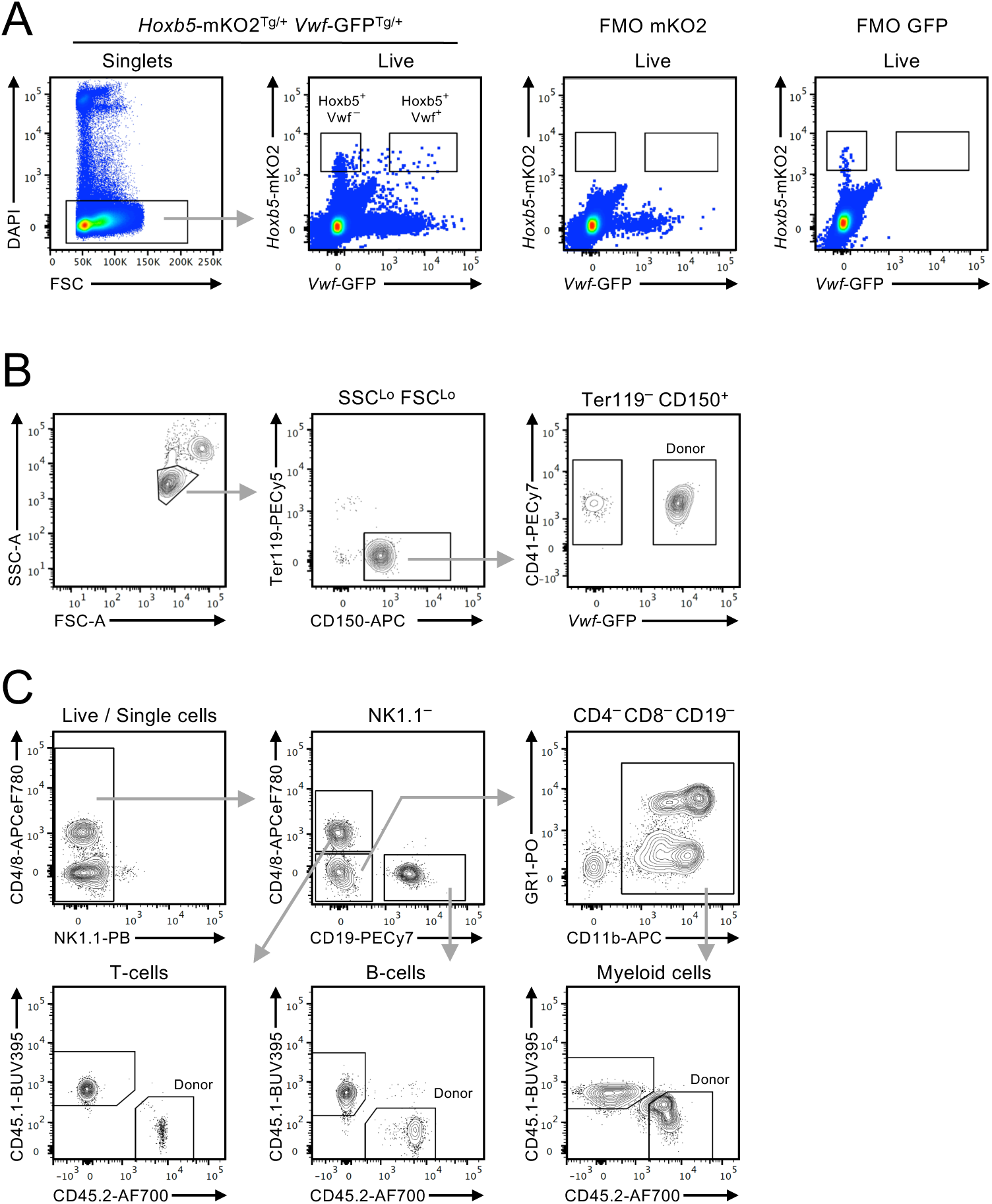
Hoxb5^+^Vwf^+^ cells have high long-term reconstitution potential. Related to main Figure 3. **A)** Sorting strategy of Hoxb5^+^Vwf^+^ and Hoxb5^+^Vwf^-^ cells from total bone marrow and solely based on the expression of the reporter transgenes. Right FACS plots show the FMO controls for mKO2 and GFP expression, used to inform the sorting gates. **B-C)** Representative FACS profiles and gating strategies for the analysis of donor chimerism in the platelets lineage **(B)** and in the myeloid, B Lymphoid and T Lymphoid cell lineages **(C)**.

**Supplementary Figure 3.**
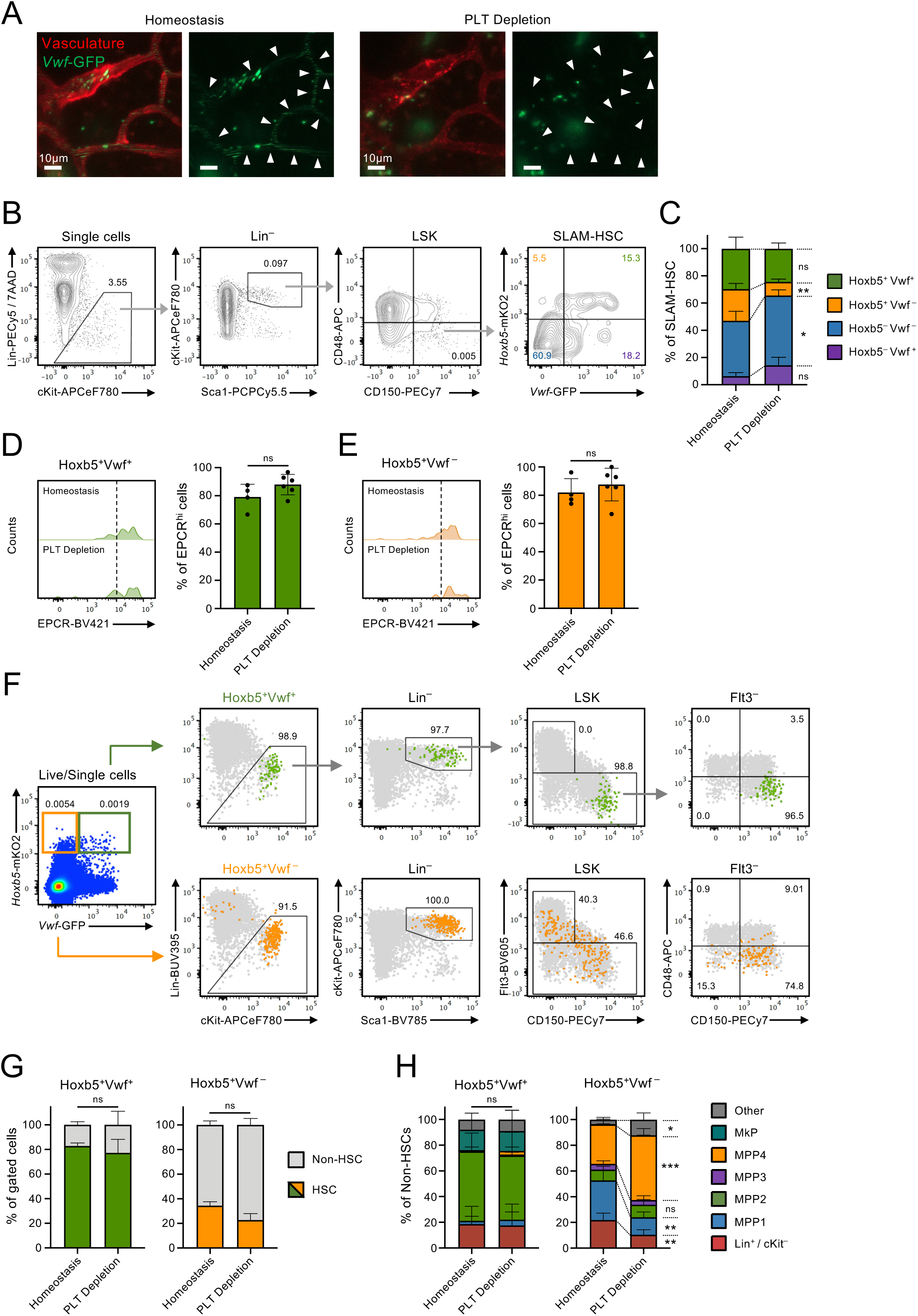
The Hoxb5^+^Vwf^+^ population is highly enriched for phenotypically defined HSCs in homeostasis and 24 hrs post platelet depletion. Related to main Figures 4-6. **A)** Representative intravital microscopy images of circulating *Vwf*-GFP^+^ platelets within calvarium blood vessels (anti-CD31/CD144-AF647 signal in red) before (homeostasis) and 2 hrs post anti-GPIbα antibody administration (PLT depletion), confirming virtually complete platelet depletion. Arrows indicate the location of platelets, that appear as lines within blood vessels of mice in homeostasis, because of their fast flow, and that are absent post platelet depletion. Scale bars represent 10 μm. **B-C)** Flow cytometric analysis of the *Hoxb5*-mKO2 and *Vwf*-GFP reporters’ expression in phenotypically defined SLAM-HSCs from mice 24 hrs post platelet depletion. **B)** Representative gating strategy of SLAM-HSCs showing the frequency of SLAM-HSC subsets defined by *Hoxb5*-mKO2 and *Vwf*-GFP expression. Numbers in gates/quadrants indicate the frequency of the gated cell populations among single cells except for the last plot (right) where they represent frequencies of the parent SLAM-HSC population, for the representative mouse shown. Data in **(C)** represents mean±SD of 4 mice in homeostasis (same mice depicted in Fig.1B) and 6 mice post platelet depletion, in 3 independent experiments. **D-E)** Flow cytometric analysis of EPCR expression within the HSC-SLAM Hoxb5^+^Vwf^+^ **(D)** and Hoxb5^+^Vwf^-^ **(E)** populations. Numbers in histograms indicate frequencies of EPCR^hi^ cells within the indicated populations. Data represents mean±SD of 4 mice in homeostasis and 6 mice 24 hrs post platelet depletion, from 3 independent experiments. **F-H)** Phenotypic characterization of total Hoxb5^+^Vwf^+^ and Hoxb5^+^Vwf^-^ cells in the BM from mice 24hrs post platelet depletion, defined exclusively based on the expression of the reporter transgenes. **F)** Hoxb5^+^Vwf^+^ (green) and Hoxb5^+^Vwf^-^ (orange) cells were gated from total BM live single cells and analysed for the frequency of SLAM-HSCs. Numbers next to gates/quadrants indicate frequency of the gated cells among the parent population. **G)** mean±SD frequency of SLAM-HSCs within the Hoxb5^+^Vwf^+^ (left) and Hoxb5^+^Vwf^-^ (right) populations. Grey cells in the background represent total BM cells undergoing same gating strategy. **H)** Phenotypic analysis of the populations represented within the non-HSC fractions of the total Hoxb5^+^Vwf^+^ (left) and Hoxb5^+^Vwf^-^ (right) cells in BM. “Other” include cells outside defined gates and populations with <0.5% representation (including pMegE, pCFU-E, pGM and GMP). **G-H**) Data are mean±SD frequencies from 8 mice in homeostasis and 8 mice post platelet depletion, in 4 independent experiments. *, p<0.05; **, p<0.01; ***, p<0.001; ns, non-significant (p>0.05); using 2-way ANOVA with Sydak’s multiple comparisons **(C, H)** or t-test **(D-E, G)**.

**Supplementary Figure 4.**
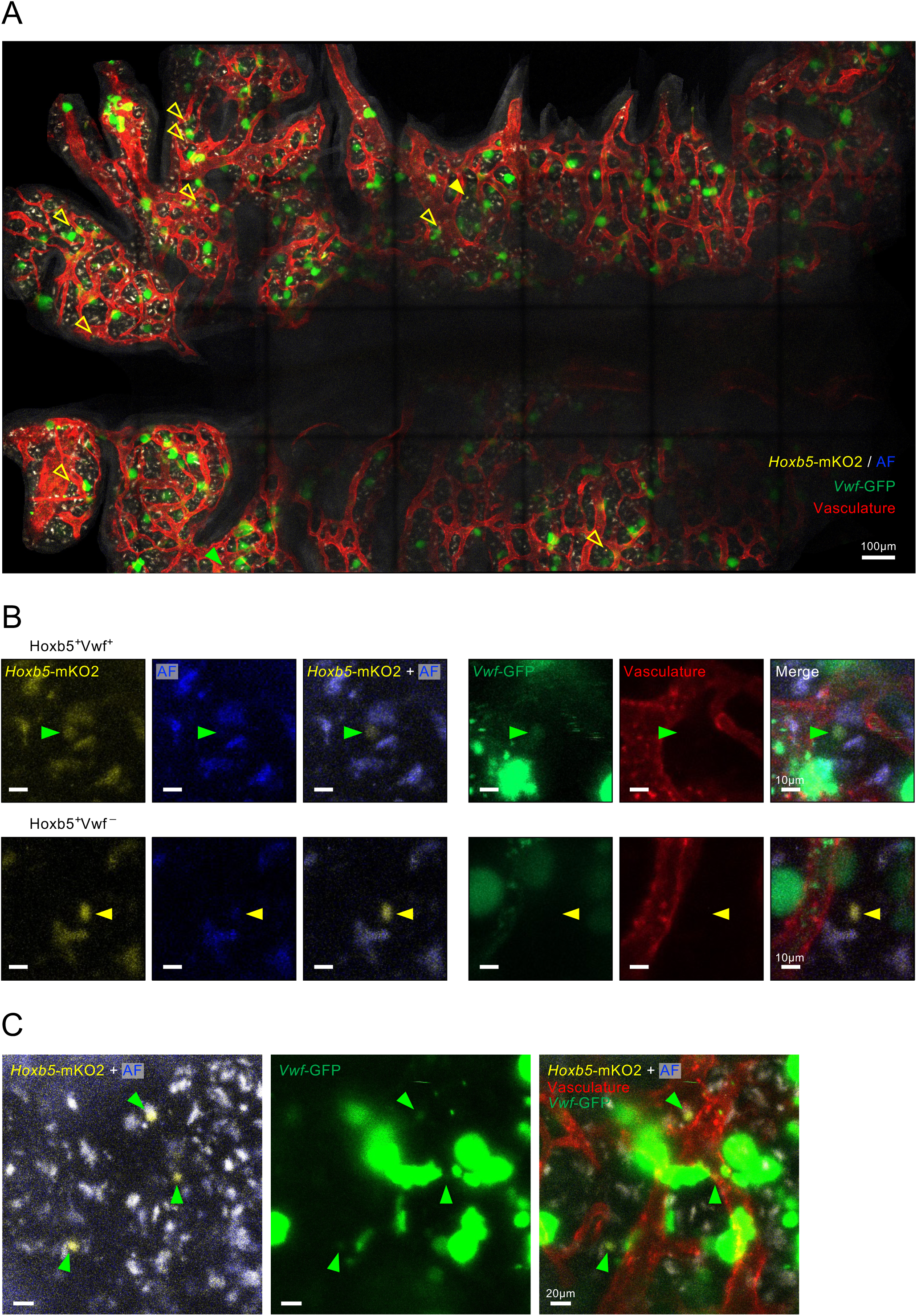
Intravital microscopy of Hoxb5-mKO2:Vwf-GFP reporter mice allows tracking of rare Hoxb5^+^Vwf^+^ cells. Related to main Figures 4 and 5. **A)** Representative maximum projection of a tilescan image of the calvarium BM of a *Hoxb5*-mKO2^+^ *Vwf*-GFP^+^ mouse in homeostasis. Red signal shows the vasculature stained by intravenous injection of AF647 conjugated anti-CD31/CD144 antibodies. Larger *Vwf-*GFP^+^ cells are Megakaryocytes. Green arrowhead in the lower part of the image indicates a Hoxb5^+^Vwf^+^ cell and yellow arrowheads indicate Hoxb5^+^Vwf^-^ cells. Filled arrowheads indicate the cells shown at higher magnification in **(B)**. AF (dim blue signal): autofluorescence. Scale bar represents 100 μm. **B)** Identification of Hoxb5^+^Vwf^+^ (top panels) and Hoxb5^+^Vwf^-^ (bottom panels) cells based on mKO2, autofluorescence (AF) and GFP signals. Scale bars represent 10 μm.

**Supplementary Figure 5.**
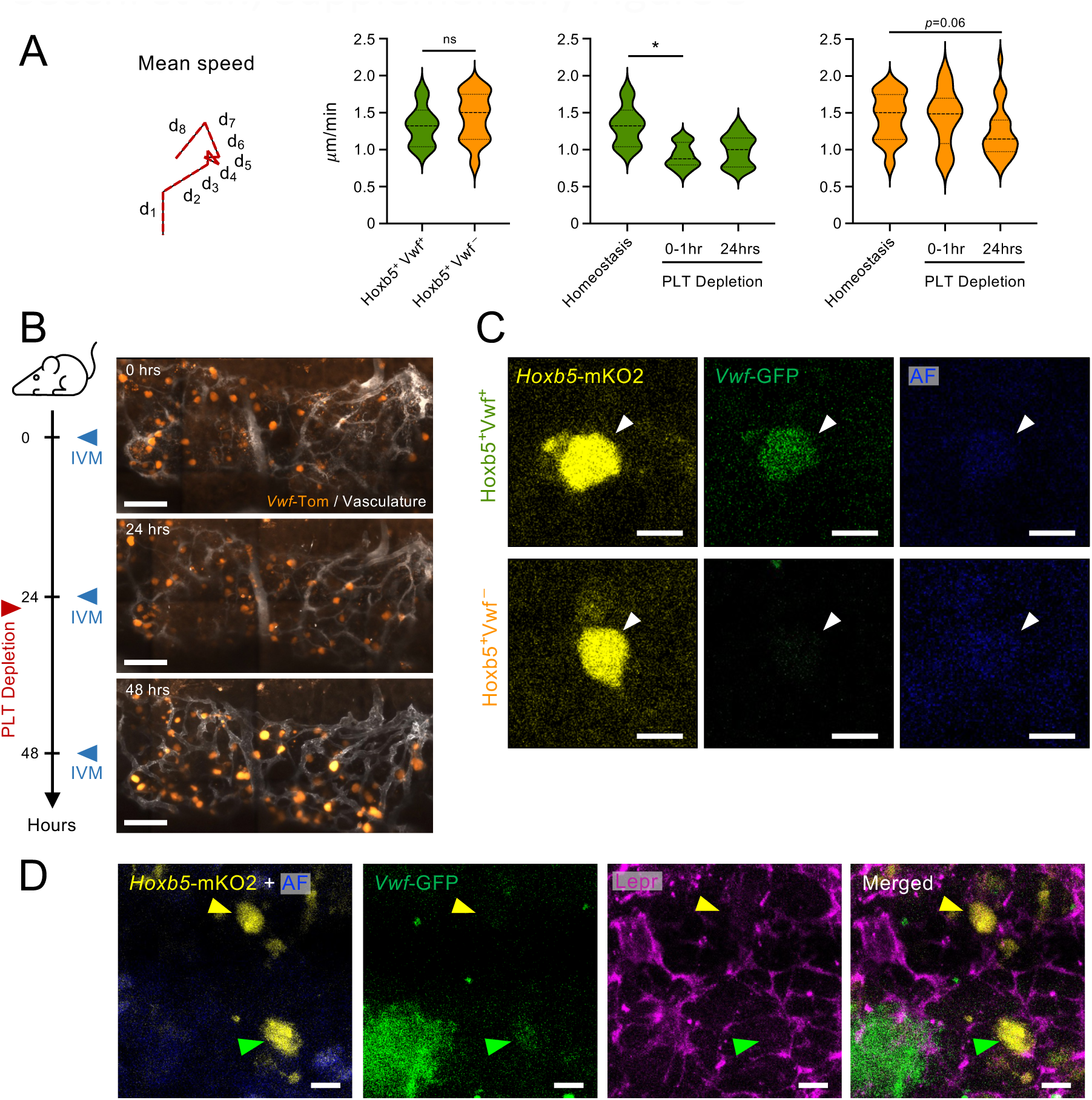
Hoxb5^+^Vwf^+^ cells localization in the bone marrow niche. Related to main Figures 5 and 6. **A)** Mean speed measurements of Hoxb5^+^Vwf^+^ and Hoxb5^+^Vwf^-^ cells in homeostasis (left graph) and of Hoxb5^+^Vwf^+^ (middle graph) or Hoxb5^+^Vwf^-^ (right graphs) cells 0-1hr post anti-GPIbα antibody administration and 24 hrs post platelet depletion. *, p<0.05; ns, non-significant (p>0.05); using Mann-Whitney test for the comparison of cells in homeostasis (left graph) or Kruskal-Wallis test with Dunn’s multiple comparisons for the comparison of cells in homeostasis versus platelet depletion (middle and right graphs). **B)** Representative images of mice imaged at the indicated time points in homeostasis and post platelet depletion. MKs were identified based on *Vwf*-tdTomato expression. Scale bars, 150 μm. **C)** Identification of Hoxb5^+^Vwf^+^ and Hoxb5^+^Vwf^-^ cells in thick section of femurs. TauSense and an empty channel channel was used to exclude autofluorescence and identify real mKO2 signal. Scale bars, 5 μm. **D)** Representative example of Hoxb5^+^Vwf^+^ and Hoxb5^+^Vwf^-^ cells in relation to LepR^+^ cells in femur bone marrow of a mouse in homeostasis. Scale bars, 5 μm.

**Supplementary Figure 6.**
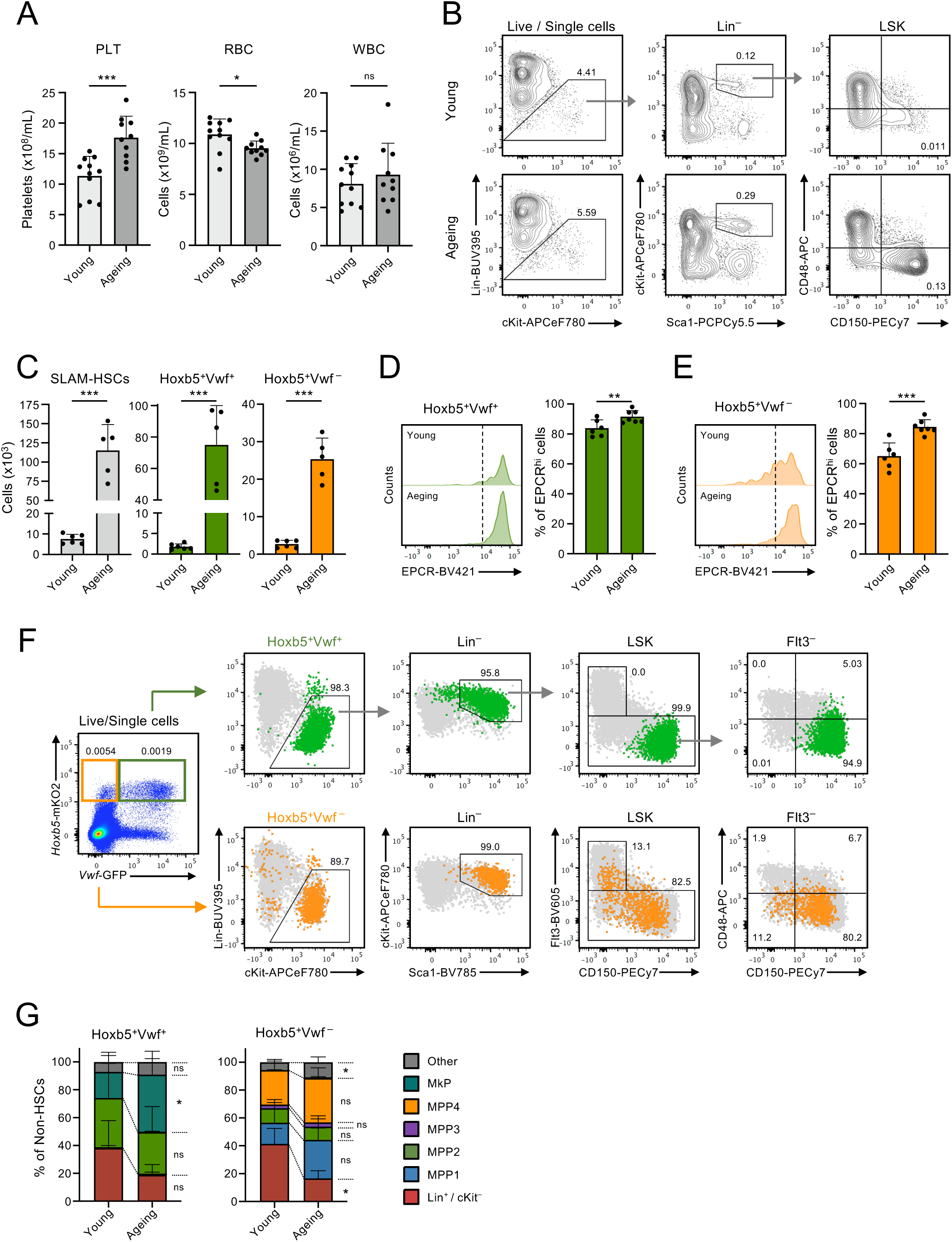
Combined expression of *Hoxb*5-mKO2 and *Vwf*-GFP defines highly enriched phenotypic HSC populations in ageing. Related to main Figure 7. **A)** Blood cell parameters of young and 1 year old mice (N=11 and 10 mice, respectively). **B-C)** Flow cytometric analysis of the *Hoxb5*-mKO2 and *Vwf*-GFP reporters’ expression in phenotypically defined SLAM-HSCs. **B)** Representative gating strategy of SLAM-HSCs and **(C)** absolute numbers of SLAM HSCs, and Hoxb5^+^Vwf^+^ and Hoxb5^+^Vwf^-^ HSC subsets from young and 1 year old mice. In **(C)** data are mean±SD from 6 young and 5 ageing mice from 3 independent experiments. **D-E)** Flow cytometric analysis of EPCR expression within Hoxb5^+^Vwf^+^ **(D)** and Hoxb5^+^Vwf^-^ **(E)** SLAM-HSC subsets from young and 1 year old mice. Data are mean±SD from 6 young and 7 ageing mice from 3 independent experiments. **F-G)** Phenotypic characterization of total Hoxb5^+^Vwf^+^ and Hoxb5^+^Vwf^-^ cells in BM, defined exclusively based on the expression of the reporter transgenes. **F)** Hoxb5^+^Vwf^+^ (green) and Hoxb5^+^Vwf^-^ (orange) cells were electronically gated from total BM live single cells and analysed for the frequency of SLAM-HSCs. Numbers next to gates/quadrants indicate frequency of the gated cells among the parent population. Grey cells in the background represent total BM cells undergoing same gating strategy. **G)** Phenotypic analysis of the populations represented within the non-HSC fraction of the total Hoxb5^+^Vwf^+^ and Hoxb5^+^Vwf^-^ cells in BM. “Other” include cells outside defined gates and populations with <0.5% representation (including pMegE, pCFU-E, pGM and GMP). Data are mean±SD frequencies from 6 young and 9 ageing mice from 4 independent experiments. *, p<0.05; **, p<0.01; ***, p<0.001; ns, non-significant (p>0.05); using t-test (**A, C-E)** or 2-way ANOVA with Sydak’s multiple comparisons **(G)**.

**Supplementary Figure 7.**
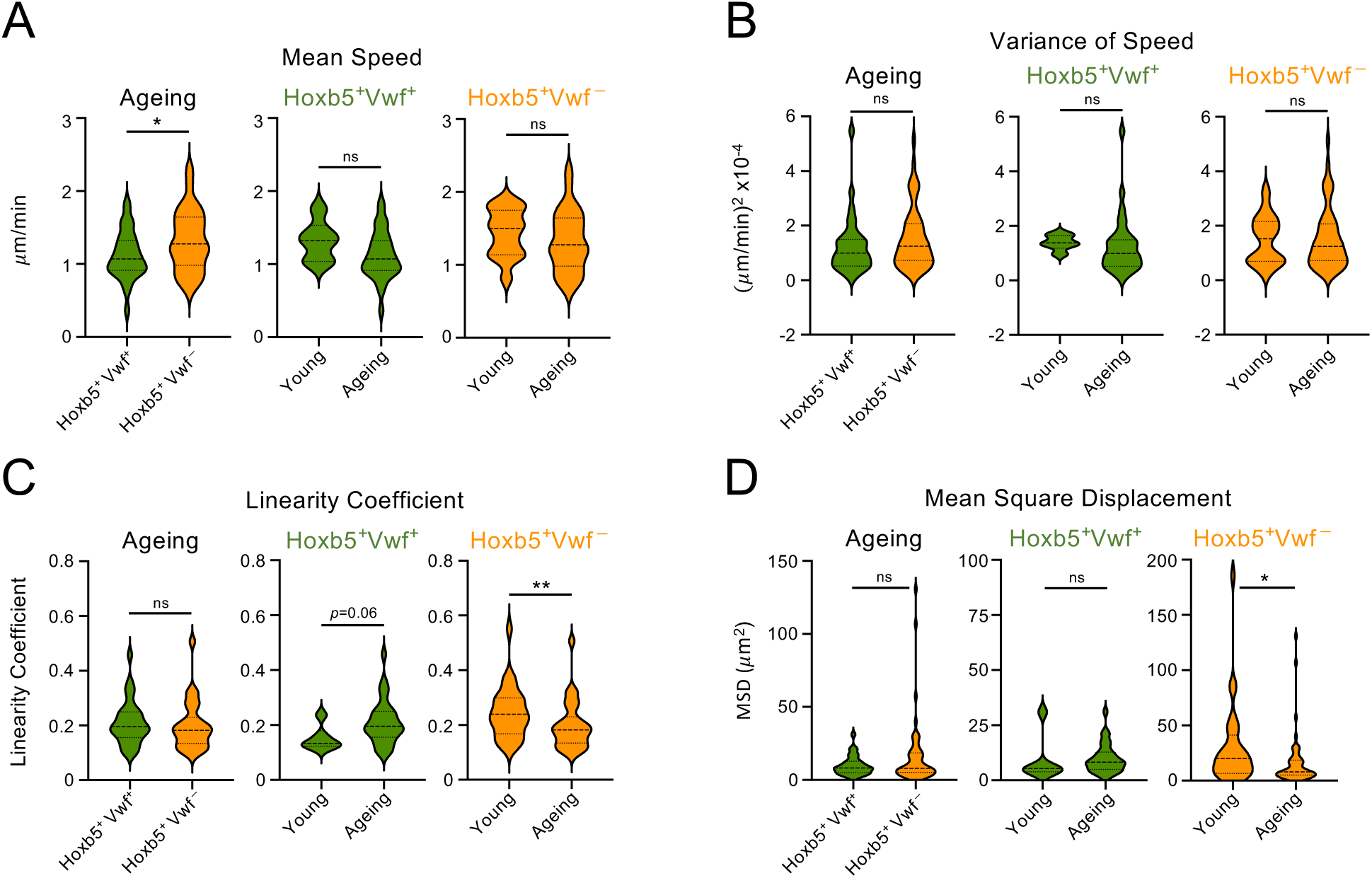
Intravital microscopy of Hoxb5^+^Vwf^+^ and Hoxb5^+^Vwf^-^ cells in calvarium BM of ageing mice. Related to main Figure 7. Track parameters extracted from 3D time-lapse datasets of single Hoxb5^+^Vwf^+^ and Hoxb5^+^Vwf^-^ cells. Violin plots depict mean speed **(A)**, mean square displacement **(B)** and linearity coefficient **(C)**. Data from 32 Hoxb5^+^Vwf^+^ cells and 56 Hoxb5^+^Vwf^-^ cells from a total of 17 mice (1 year old) imaged. Data depicted for young mice are the same shown in Figures 3 and 4, corresponding to 6 Hoxb5^+^Vwf^+^ cells and 28 Hoxb5^+^Vwf^-^ cells from 12 mice in homeostasis. *, p<0.05; **, p<0.01; ns, non-significant (p>0.05); using Mann-Whitney test.

**Supplementary Table 1.** Flow Cytometry antibodies.

| Antigen | Conjugate | Clone | Supplier | Catalogue No. | Dilution |
| --- | --- | --- | --- | --- | --- |
| B220 | BUV395 | RA3-6B2 | BD | 563793 | 1/200 |
| B220 | PECy5 | RA3-6B2 | BioLegend | 103210 | 1/400 |
| CD105 | BV421 | MJ7/18 | BD | 562760 | 1/100 |
| CD117 (cKit) | APCeF780 | 2B8 | Invitrogen | 47-1171-82 | 1/1200 |
| CD11b | APC | M1/70 | BioLegend | 101212 | 1/600 |
| CD135 (Flt3) | Bio | A2F10 | Invitrogen | 13-1351-85 | 1/100 |
| CD150 | APC | TC15-12F12.2 | BioLegend | 115910 | 1/100 |
| CD150 | PECy7 | TC15-12F12.2 | BioLegend | 115914 | 1/400 |
| CD16/32 (FcγRII/III) | AF700 | 93 | Invitrogen | 56-0161-80 | 1/100 |
| CD19 | PECy7 | ebio1D3 | Invitrogen | 25-0193-82 | 1/400 |
| CD201 (EPCR) | Bio | ebio1560 | Invitrogen | 13-2012-82 | 1/200 |
| CD34 | AF700 | RAM34 | Invitrogen | 56-0341-82 | 1/10 |
| CD4 | APCeF780 | RM4-5 | Invitrogen | 47-0042-82 | 1/1600 |
| CD4 | BUV395 | RM4-5 | BD | 740208 | 1/400 |
| CD4 | PECy5 | RM4-5 | BioLegend | 100514 | 1/800 |
| CD41 | BUV737 | MWReg30 | BD | 741759 | 1/300 |
| CD41 | PECy7 | eBioMWReg30 | Invitrogen | 25-0411-82 | 1/800 |
| CD45.1 | BUV395 | A20 | BD | 565212 | 1/200 |
| CD45.2 | AF700 | 104 | BioLegend | 109822 | 1/50 |
| CD48 | APC | HM48-1 | BioLegend | 103412 | 1/600 |
| CD5 | BUV395 | 53-7.3 | BD | 740206 | 1/100 |
| CD5 | PECy5 | 53-7.3 | BioLegend | 100610 | 1/800 |
| CD8 | APCeF780 | 53-6.7 | Invitrogen | 47-0081-82 | 1/800 |
| CD8 | BUV395 | 53-6.7 | BD | 563786 | 1/200 |
| CD8 | PECy5 | 53-6.7 | BioLegend | 100710 | 1/800 |
| GR1 | BUV395 | RB6-8C5 | BD | 563849 | 1/400 |
| GR1 | PECy5 | RB6-8C5 | BioLegend | 108410 | 1/800 |
| GR1 | PO | RB6-8C5 | Invitrogen | RM3030 | 1/100 |
| NK1.1 | PB | PK136 | BioLegend | 108722 | 1/800 |
| SAv | BV421 | - | BioLegend | 405226 | 1/200 |
| SAv | BV605 | - | BioLegend | 405229 | 1/200 |
| Sca1 | BV785 | D7 | BioLegend | 108139 | 1/200 |
| Sca1 | PCPCy5.5 | D7 | Invitrogen | 45-5981-80 | 1/300 |
| Ter119 | BUV395 | TER-119 | BD | 563827 | 1/100 |
| Ter119 | PECy5 | TER-119 | BioLegend | 116210 | 1/600 |

**Supplementary Table 2.** Imaging antibodies.

| Antigen | Conjugate | Clone | Supplier | Catalogue No. | Amount / Dilution |
| --- | --- | --- | --- | --- | --- |
| IVM |  |  |  |  |  |
| CD144 | AF647 | BV13 | BioLegend | 138006 | 10 ug |
| CD31 | AF647 | MEC13.3 | BioLegend | 102516 | 10 ug |
| Bone marrow sections |  |  |  |  |  |
| LepR | Bio | Polyclonal Goat IgG | R&D Systems | BAF497 | 1/20 |
| Anti-Goat IgG | AF800 | Polyclonal Donkey IgG | Invitrogen | A32930 | 1/250 |

## Supplementary Movies

**Supplementary Movie 1. Time-lapse IVM of Hoxb5^+^Vwf^+^ cell in homeostasis.** Related to main figure 4C.

**Supplementary Movie 2. Time-lapse IVM of Hoxb5^+^Vwf^-^ cell in homeostasis.** Related to main figure 4C.

**Supplementary Movie 3. Time-lapse IVM of Hoxb5^+^Vwf^+^ cell 0-1 hr post acute platelet depletion.** Related to main figure 4D.

**Supplementary Movie 4. Time-lapse IVM of Hoxb5^+^Vwf^-^ cell 0-1 hr post acute platelet depletion.** White arrowhead indicates the Hoxb5^+^Vwf^-^ cell. Green arrow heads indicate MKs disintegrating into pro-platelets. Related to main figure 4D.

**Supplementary Movie 5. Time-lapse IVM of Hoxb5^+^Vwf^+^ cell 24 hrs post acute platelet depletion.** Related to main figure 4E.

**Supplementary Movie 6. Time-lapse IVM of Hoxb5^+^Vwf^-^ cell 24 hrs post acute platelet depletion.** Related to main figure 4E.

**Supplementary Movie 7. Time-lapse IVM of Hoxb5^+^Vwf^+^ cell in ageing.** Related to main figure 7C.

**Supplementary Movie 8. Time-lapse IVM of Hoxb5^+^Vwf^-^ cells in ageing.** Each arrowhead depicts a different cell. Related to main figure 7C.

## REFERENCES

1 Laurenti, E. & Gottgens, B. From haematopoietic stem cells to complex differentiation landscapes. Nature 553, 418–426, doi:10.1038/nature25022 (2018).

2 Morrison, S. J. & Scadden, D. T. The bone marrow niche for haematopoietic stem cells. Nature 505, 327–334, doi:nature12984 [pii] 10.1038/nature12984 (2014).

3 Pinho, S. & Frenette, P. S. Haematopoietic stem cell activity and interactions with the niche. Nat Rev Mol Cell Biol 20, 303–320, doi:10.1038/s41580-019-0103-9 (2019).

4 Kokkaliaris, K. D. et al. Adult blood stem cell localization reflects the abundance of reported bone marrow niche cell types and their combinations. Blood 136, 2296–2307, doi:10.1182/blood.2020006574 (2020).

5 Baccin, C. et al. Combined single-cell and spatial transcriptomics reveal the molecular, cellular and spatial bone marrow niche organization. Nat Cell Biol 22, 38–48, doi:10.1038/s41556-019-0439-6 (2020).

6 Baryawno, N. et al. A Cellular Taxonomy of the Bone Marrow Stroma in Homeostasis and Leukemia. Cell 177, 1915–1932 e1916, doi:10.1016/j.cell.2019.04.040 (2019).

7 Tikhonova, A. N. et al. The bone marrow microenvironment at single-cell resolution. Nature 569, 222–228, doi:10.1038/s41586-019-1104-8 (2019).

8 Acar, M. et al. Deep imaging of bone marrow shows non-dividing stem cells are mainly perisinusoidal. Nature 526, 126–130, doi:10.1038/nature15250 (2015).

9 Chen, J. Y. et al. Hoxb5 marks long-term haematopoietic stem cells and reveals a homogenous perivascular niche. Nature 530, 223–227, doi:10.1038/nature16943 (2016).

10 Christodoulou, C. et al. Live-animal imaging of native haematopoietic stem and progenitor cells. Nature 578, 278–283, doi:10.1038/s41586-020-1971-z (2020).

11 Gazit, R. et al. Fgd5 identifies hematopoietic stem cells in the murine bone marrow. J Exp Med 211, 1315–1331, doi:10.1084/jem.20130428 (2014).

12 Kucinski, I. et al. A time- and single-cell-resolved model of murine bone marrow hematopoiesis. Cell Stem Cell 31, 244–259 e210, doi:10.1016/j.stem.2023.12.001 (2024).

13 Nakatani, T. et al. Ebf3(+) niche-derived CXCL12 is required for the localization and maintenance of hematopoietic stem cells. Nat Commun 14, 6402, doi:10.1038/s41467-023-42047-2 (2023).

14 Rodriguez-Fraticelli, A. E. et al. Single-cell lineage tracing unveils a role for TCF15 in haematopoiesis. Nature 583, 585–589, doi:10.1038/s41586-020-2503-6 (2020).

15 Takihara, Y. et al. Bone marrow imaging reveals the migration dynamics of neonatal hematopoietic stem cells. Commun Biol 5, 776, doi:10.1038/s42003-022-03733-x (2022).

16 Upadhaya, S. et al. Intravital Imaging Reveals Motility of Adult Hematopoietic Stem Cells in the Bone Marrow Niche. Cell Stem Cell 27, 336–345 e334, doi:10.1016/j.stem.2020.06.003 (2020).

17 Haltalli, M. L. R. & Lo Celso, C. Intravital Imaging of Bone Marrow Niches. Methods Mol Biol 2308 203–222, doi:10.1007/978-1-0716-1425-9_16 (2021).

18 Foster, K. et al. Different Motile Behaviors of Human Hematopoietic Stem versus Progenitor Cells at the Osteoblastic Niche. Stem Cell Reports 5, 690–701, doi:10.1016/j.stemcr.2015.09.003 (2015).

19 Lo Celso, C., et al. Live-animal tracking of individual haematopoietic stem/progenitor cells in their niche. Nature 457, 92–96, doi:nature07434 [pii] 10.1038/nature07434 (2009).

20 Rashidi, N. M. et al. In vivo time-lapse imaging of mouse bone marrow reveals differential niche engagement by quiescent and naturally activated hematopoietic stem cells. Blood, doi:blood-2013-10-534859 [pii] 10.1182/blood-2013-10-534859 (2014).

21 Challen, G. A., Boles, N. C., Chambers, S. M. & Goodell, M. A. Distinct hematopoietic stem cell subtypes are differentially regulated by TGF-beta1. Cell Stem Cell 6, 265–278, doi:S1934-5909(10)00046-9 [pii] 10.1016/j.stem.2010.02.002 (2010).

22 Dykstra, B. et al. Long-term propagation of distinct hematopoietic differentiation programs in vivo. Cell Stem Cell 1, 218–229, doi:S1934-5909(07)00021-5 [pii] 10.1016/j.stem.2007.05.015 (2007).

23 Muller-Sieburg, C. E., Cho, R. H., Thoman, M., Adkins, B. & Sieburg, H. B. Deterministic regulation of hematopoietic stem cell self-renewal and differentiation. Blood 100, 1302–1309 (2002).

24 Carrelha, J. et al. Hierarchically related lineage-restricted fates of multipotent haematopoietic stem cells. Nature 554, 106–111, doi:10.1038/nature25455 (2018).

25 Rodriguez-Fraticelli, A. E. et al. Clonal analysis of lineage fate in native haematopoiesis. Nature 553, 212–216, doi:10.1038/nature25168 (2018).

26 Sanjuan-Pla, A. et al. Platelet-biased stem cells reside at the apex of the haematopoietic stem-cell hierarchy. Nature 502, 232–236, doi:10.1038/nature12495 (2013).

27 Yamamoto, R. et al. Clonal analysis unveils self-renewing lineage-restricted progenitors generated directly from hematopoietic stem cells. Cell 154, 1112–1126, doi:10.1016/j.cell.2013.08.007 (2013).

28 Carrelha, J. et al. Alternative platelet differentiation pathways initiated by nonhierarchically related hematopoietic stem cells. Nat Immunol 25, 1007–1019, doi:10.1038/s41590-024-01845-6 (2024).

29 Luis, T. C. et al. Perivascular niche cells sense thrombocytopenia and activate hematopoietic stem cells in an IL-1 dependent manner. Nat Commun 14, 6062, doi:10.1038/s41467-023-41691-y (2023).

30 Bruns, I. et al. Megakaryocytes regulate hematopoietic stem cell quiescence through CXCL4 secretion. Nat Med 20, 1315–1320, doi:10.1038/nm.3707 (2014).

31 Zhao, M. et al. Megakaryocytes maintain homeostatic quiescence and promote post-injury regeneration of hematopoietic stem cells. Nat Med 20, 1321–1326, doi:10.1038/nm.3706 (2014).

32 Pinho, S. et al. Lineage-Biased Hematopoietic Stem Cells Are Regulated by Distinct Niches. Dev Cell 44, 634–641 e634, doi:10.1016/j.devcel.2018.01.016 (2018).

33 Kunisaki, Y. et al. Arteriolar niches maintain haematopoietic stem cell quiescence. Nature 502, 637–643, doi:nature12612 [pii] 10.1038/nature12612 (2013).

34 Florez, M. A. et al. Interferon Gamma Mediates Hematopoietic Stem Cell Activation and Niche Relocalization through BST2. Cell Rep 33, 108530, doi:10.1016/j.celrep.2020.108530 (2020).

35 Nestorowa, S. et al. A single-cell resolution map of mouse hematopoietic stem and progenitor cell differentiation. Blood 128, e20–31, doi:10.1182/blood-2016-05-716480 (2016).

36 Hamey, F. K. & Gottgens, B. Machine learning predicts putative hematopoietic stem cells within large single-cell transcriptomics data sets. Exp Hematol 78, 11–20, doi:10.1016/j.exphem.2019.08.009 (2019).

37 Wilson, N. K. et al. Combined Single-Cell Functional and Gene Expression Analysis Resolves Heterogeneity within Stem Cell Populations. Cell Stem Cell 16, 712–724, doi:10.1016/j.stem.2015.04.004 (2015).

38 Che, J. L. C. et al. Identification and characterization of in vitro expanded hematopoietic stem cells. EMBO Rep 23, e55502, doi:10.15252/embr.202255502 (2022).

39 Grover, A. et al. Single-cell RNA sequencing reveals molecular and functional platelet bias of aged haematopoietic stem cells. Nat Commun 7, 11075, doi:10.1038/ncomms11075 (2016).

40 Young, K. et al. Decline in IGF1 in the bone marrow microenvironment initiates hematopoietic stem cell aging. Cell Stem Cell 28, 1473–1482 e1477, doi:10.1016/j.stem.2021.03.017 (2021).

41 Lin, D. S. & Trumpp, A. Differential expression of endothelial protein C receptor (EPCR) in hematopoietic stem and multipotent progenitor cells in young and old mice. Cells Dev 174, 203843, doi:10.1016/j.cdev.2023.203843 (2023).

42 Sanchez-Aguilera, A. et al. Estrogen signaling selectively induces apoptosis of hematopoietic progenitors and myeloid neoplasms without harming steady-state hematopoiesis. Cell Stem Cell 15, 791–804, doi:10.1016/j.stem.2014.11.002 (2014).

43 Li, J. J. et al. Differentiation route determines the functional outputs of adult megakaryopoiesis. Immunity 57, 478–494 e476, doi:10.1016/j.immuni.2024.02.006 (2024).

44 Morcos, M. N. F. et al. Fate mapping of hematopoietic stem cells reveals two pathways of native thrombopoiesis. Nat Commun 13, 4504, doi:10.1038/s41467-022-31914-z (2022).

45 Poscablo, D. M. et al. An age-progressive platelet differentiation path from hematopoietic stem cells causes exacerbated thrombosis. Cell 187, 3090–3107 e3021, doi:10.1016/j.cell.2024.04.018 (2024).

46 Nakamura-Ishizu, A., Takubo, K., Fujioka, M. & Suda, T. Megakaryocytes are essential for HSC quiescence through the production of thrombopoietin. Biochem Biophys Res Commun 454, 353–357, doi:10.1016/j.bbrc.2014.10.095 (2014).

47. Pylvanainen, J. W., et al. Fast4DReg - fast registration of 4D microscopy datasets. J Cell Sci 136, doi:10.1242/jcs.260728 (2023).

48 Beltman, J. B., Maree, A. F. & de Boer, R. J. Analysing immune cell migration. Nat Rev Immunol 9, 789–798, doi:10.1038/nri2638 (2009).

49 Wolf, F. A., Angerer, P. & Theis, F. J. SCANPY: large-scale single-cell gene expression data analysis. Genome Biol 19, 15, doi:10.1186/s13059-017-1382-0 (2018).

50 Dobin, A. et al. STAR: ultrafast universal RNA-seq aligner. Bioinformatics 29, 15–21, doi:10.1093/bioinformatics/bts635 (2013).

51 Liao, Y., Smyth, G. K. & Shi, W. featureCounts: an efficient general purpose program for assigning sequence reads to genomic features. Bioinformatics 30, 923–930, doi:10.1093/bioinformatics/btt656 (2014).

